# Generation of antigen-specific and functionally stable Treg cells from effector/memory T cells for cell therapy of immunological diseases

**DOI:** 10.1101/2025.06.13.659621

**Authors:** Norihisa Mikami, Ryoji Kawakami, Atsushi Sugimoto, Masaya Arai, Shimon Sakaguchi

## Abstract

One strategy for antigen-specific immunosuppression is to convert antigen-specific conventional T cells (Tconvs) into Foxp3^+^ regulatory T cells (Tregs) that are as stably suppressive as naturally occurring Tregs (nTregs). To achieve the conversion *in vitro* for mice and humans, we induced high Foxp3 expression in antigen- and IL-2-stimulated Tconvs by CDK8/19 inhibition, and established Treg-specific epigenetic changes by depriving CD28 co-stimulation during *in vitro* Treg induction to specifically promote the expression of Treg signature genes, especially *Foxp3*. Repeating this process, with intermittent resting cultures containing IL-2 only, enabled in a short duration efficient conversion of naïve as well as effector/memory CD4^+^ Tconvs, including Th1, Th2 and Th17 cells, into Foxp3^+^ Tregs, which were similar to nTregs in transcription and epigenetic modification. Induced Tregs (iTregs) thus generated were indeed functionally and phenotypically stable *in vivo* and effectively suppressed inflammatory bowel disease and graft-versus-host disease in mouse models. Adoptive cell therapy with such effector/memory Tconv-derived, functionally stable, iTregs would be able to achieve antigen- and disease-specific treatment of immunological diseases.

**One Sentence Summary:** Antigen-specific and functionally stable Treg cells can be generated *in vitro* from effector/memory T cells for cell therapy of immunological diseases.

## Introduction

A prevailing challenge in treating autoimmune and other inflammatory diseases is how to establish stable immune suppression and reestablish self-tolerance in an antigen-specific or a disease-specific manner. There are accumulating demonstrations in animal models that regulatory T cells (Tregs) specifically expressing the transcription factor (TF) Foxp3 in the nucleus, CD25 and CTLA-4 on the cell surface, are instrumental in immune suppression and tolerance induction (1–7). Tregs are not only naturally present in the immune system (natural Tregs: nTregs); they can also be induced from conventional T cells (Tconvs) *in vitro* (induced Tregs: iTregs) (8, 9). For example, antigen stimulation of naïve CD4^+^ Tconvs in the presence of TGF-β and IL-2 is able to convert them into antigen-specific iTregs (10, 11). However, in contrast to nTregs, which are functionally stable *in vivo* and *in vitro*, iTregs are unstable especially in inflammatory environments and difficult to generate from effector/memory Tconvs (12–15). Nonetheless, antigen-specific iTregs are easier to prepare *in vitro* in large quantities as opposed to nTregs, which are resistant to *in vitro* expansion by antigenic stimulation (16, 17). These findings raise the question of how antigen-specific iTregs as functionally stable as nTregs can be produced in sufficient amounts not only from naïve Tconvs but also from those antigen-primed effector/memory Tconvs that are the actual culprits behind immunological diseases.

The establishment of both the Foxp3-dependent TF network and the Treg-specific epigenetic landscape coordinates Treg development and function (18). Foxp3, which is responsible for conferring immunosuppressive functions on Tregs, represses IL-2 and other proinflammatory cytokines and enhances the expression of Treg signature genes such as *Il2ra*, *Ctla4*, and *Foxp3* itself (3–5). Moreover, epigenetic alterations (e.g., DNA hypomethylation, histone modifications, and chromatin remodeling) especially at enhancer regions of such Treg signature genes are required for their specific and sustained expression (13, 19–22). The functional instability of iTregs may be attributed to a lack of Treg-specific epigenetic changes, especially DNA hypomethylation (12, 13). It has also been shown that Foxp3 protein expression itself hardly controls the formation of such Treg-specific epigenome in developing Tregs. For example, developing nTregs in the thymus acquire Treg-specific epigenetic changes without Foxp3 expression (13, 21, 22); retroviral ectopic expression of Foxp3 in Tconvs is unable to induce Treg-specific DNA hypomethylation in *Foxp3* and other Treg signature genes (12, 13); and such Treg-specific DNA hypomethylation can indeed enhance, independently of Foxp3, the expression of Treg signature genes including *Il2ra* and *Ctla4* in developing nTregs (13, 23). These findings collectively suggest that Foxp3 expression and the formation of Treg-type epigenome are operationally distinguishable and can be separately installed in iTregs to endow them with stable suppressive function.

TGF-β and several other small molecules such as rapamycin, retinoic acid (RA) and short-chain fatty acids (e.g., butyrate) are able to induce Foxp3, though all the latter substances require TGF-β for initial Foxp3 expression (24–26). Such TGF-β-dependent Foxp3 induction is antagonized by inflammatory cytokines such as IL-4 and IL-6, which promote IL-9-producing Th9 or IL-17-producing Th17 cells, respectively, in the presence of TGF-β (27–29). For more efficient induction of Foxp3, inhibitors of cyclin-dependent kinases (CDK) 8/19 can be used as they induce Foxp3 expression not only through a TGFβ-independent and STAT5-dependent mechanism but also block Tconv differentiation into effector Th cells in the presence of pro-inflammatory cytokines (30). As for the induction of Treg-type epigenome, deprivation of CD28 co-stimulatory signal during iTreg generation can establish Treg-specific DNA hypomethylation at enhancer regions of Treg signature genes, typically the *Foxp3* CNS2 region, even in Foxp3-deficient CD4^+^ T cells (31). Thus, it may be beneficial to combine these distinct procedures for Foxp3 induction and for Treg-specific epigenetic changes in order to generate functionally stable antigen-specific iTregs even from effector/memory Tconvs.

Here we have addressed in mice and humans how a combination of these *in vitro* measures for Foxp3 induction and Treg-specific epigenetic alterations can convert Tconvs, especially antigen-specific effector/memory Tconvs such as Th1, Th2 and Th17 effector T cells, into potently suppressive and functionally stable iTregs in sufficient numbers within a short period. The results would help devising adoptive cell therapy with antigen-specific and functionally stable iTregs for long-term immune suppression and tolerance induction in inflammatory tissue environments, enabling treatment of autoimmune and other immunological diseases and prevention of graft rejection in organ transplantation, without general immune suppression.

## Results

### Generation of functionally stable iTregs from naïve Tconvs

To generate iTregs with high Foxp3 expression and Treg-specific epigenetic alterations from lymph node naïve (CD44^lo^CD62L^hi^) CD4^+^ T cells in Foxp3-eGFP (eFox) reporter mice (**Fig. S1A**), we stimulated the cells for 3 days with plate-bound anti-CD3 mAb, IL-2 and TGF-β, with or without soluble agonistic anti-CD28 mAb, and with or without the CDK8/19 inhibitor Senexin A (First Stimulation) (**Fig. 1A**). Compared to conventional iTregs (C-iTregs), which were generated with anti-CD3 mAb, anti-CD28 mAb, TGF-β, and IL-2, the addition of Senexin A or the removal of anti-CD28 mAb, and, in particular, both treatments together, enhanced the generation of Foxp3^+^ cells. They highly expressed Foxp3, CD25, and especially CTLA-4, which is essential for Treg suppressive function to reduce CD80/CD86 and increase free PD-L1 on antigen-presenting cells (APCs) (6, 32, 33) (**Fig. 1B, 1C** and **S1B**). Since restimulation after the first stimulation rendered the cells prone to die by activation-induced cell death with high expression of Fas-ligand (FasL) (**Fig. S1C**) (34), we cultured the cells with IL-2 alone for 4 days (Resting Culture), then restimulated them in a similar manner as the first stimulation for 2 more days, with addition of anti-FasL mAb to prevent cell death (Second Stimulation) (**Fig. 1A** and **S1C**). Notably, in the second stimulation, the combination of Senexin A and the removal of anti-CD28 mAb further enhanced the ratio of Foxp3^+^ cells to nearly 100%, with higher levels of Foxp3, CD25, and CTLA-4 expression than the first stimulation (**Fig. 1B, 1C** and **S1B**). They had stable Foxp3 expression in extended *in vitro* culture even in the presence of inflammatory cytokines (**Fig. S1D**). Moreover, this protocol with the secondary stimulation generated Foxp3^+^ cells more efficiently compared to the other protocols, yielding ∼2×10^7^ Foxp3^+^ cells from 10^5^ Foxp3^-^ naïve CD4^+^ T cells in 10 days (**Fig. 1D**). Hereafter, we call iTregs produced by this protocol as stable and functional iTregs (S/F-iTregs).

**Fig. 1.**
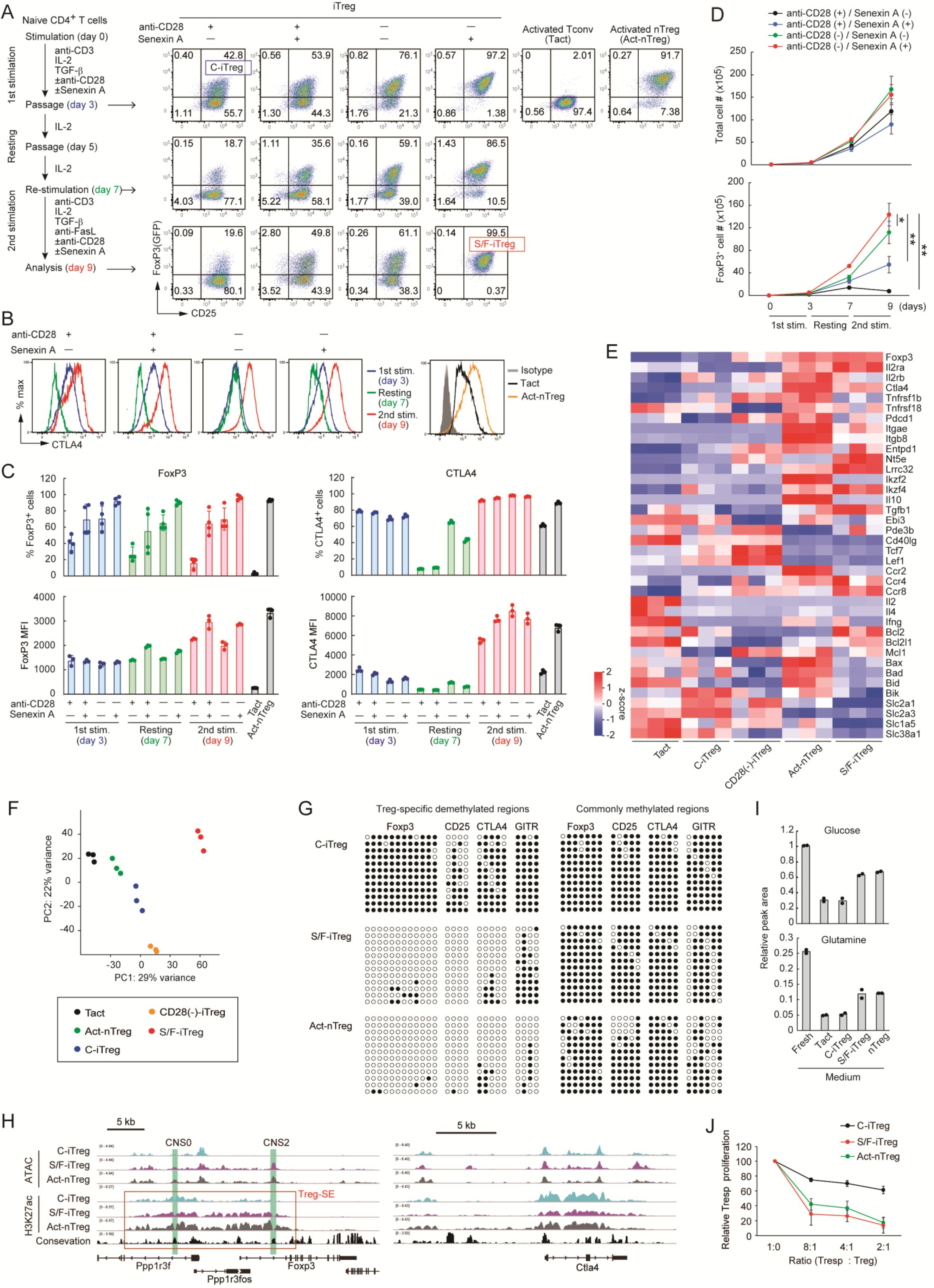
Generation of stable and functional iTregs from naïve CD4^+^ Tconvs. (**A**) Protocol for iTreg induction from naïve (CD44^lo^CD62L^hi^) CD4^+^ Tconv suspensions from LNs of normal mice and Foxp3/CD25 staining of CD4^+^ T cells at each step of iTreg induction. Representative of four independent experiments. (**B**) CTLA-4 staining of CD4^+^ Tconvs at each step of iTreg induction shown in (A). (**C**) Percentages of Foxp3^+^ and CTLA-4^+^ cells, their Foxp3 and CTLA-4 MFIs, assessed by flow cytometry at each step of iTreg induction. (**D**) The numbers of total and FoxP3^+^ cells at each step of iTreg induction. Vertical bars denote mean±SD (n=3-4) in C and D. (**E, F**) Gene expression pattern (E) and PCA (F) of Treg-related molecules in *in vitro* activated Tconvs (Tacts), conventional iTregs (C-iTregs), iTregs prepared without agonistic anti-CD28 mAb (CD28(-)-iTregs), *in vitro* stimulated nTregs, and stable/functional iTregs (S/F-iTregs), as show in Fig. 1A (n=3). (**G**) DNA hypomethylation of Treg-specific demethylated regions and constitutively methylated regions assessed by bisulfite sequencing analysis. Filled circle; methylated CpG, unfilled circle; demethylated CpG. (**H**) Chromatin accessibility and enhancer activity assessed by ATAC-seq and H3K27ac ChIP-seq, respectively. Red square denotes Treg-specific super-enhancer (Treg-SE) region. Representative of 2 independent experiments in G and H. (**I**) Relative amount of glucose and glutamine conditional medium of each T cell subset measured by mass spectrometry (two biological replicates). (**J**) Suppressive activity of Treg populations. Proliferation of Treg-cocultured CFSE-labelled responder CD4^+^ T cells (Tresp) in the presence of anti-CD3 and APCs was assessed and expressed as percentages compared with proliferation of Tresp cells alone as 100%. Vertical bars denote mean±SD (n=3). ANOVA followed by SNK test for statistical analysis in D (* p<0.05, ** p<0.01).

S/F-iTregs exhibited a Treg-specific gene expression profile, as assessed by RNA-seq, that was more representative of activated nTregs than C-iTregs (**Fig. 1E**). They were distinct from the first-stimulation iTregs by PCA analysis, indicating that the major transcriptome alterations in S/F-iTregs could be attributed to the second stimulation (**Fig. 1F** and **S1E**). For example, *Tgfb1* and *Lrrc32* (encoding GARP) were highly transcribed, and the proteins highly expressed, in both S/F-iTregs and nTregs compared to C-iTregs (**Fig. 1E** and **S1F**), while *Cd40lg*, *Tcf7*, and *Lef1* were similarly low in both S/F-iTreg and nTreg populations (**Fig. 1E**). Also, S/F-iTregs and activated nTregs had some noticeable differences; for example, the former were lower in the expression of *Ikzf2* (Helios) and IL-10 at the gene and protein levels (**Fig. S1G** and below), and higher in anti-apoptotic *Bcl2*. Pro-apoptotic genes such as *Bax*, *Bad* and *Bid* were also lower in S/F-iTregs, presumably due to the Fas-Ligand blockade (**Fig. S1H**). Assessment of epigenetic statuses of Treg-signature genes, such as *Foxp3*, *Il2ra*, *Ctla4*, and *Tnfrsf18,* revealed that the enhancer regions of these genes (e.g., the CNS2 region of *Foxp3*) in S/F-iTregs possessed specific DNA hypomethylation as present in nTregs but not in C-iTregs (13) (**Fig. 1G**). S/F-iTregs were also comparable to nTregs in high chromatin accessibility and enhancer activation at the *Foxp3* and *Ctla4* loci, as assessed by ATAC-seq and H3K27ac ChIP-seq, respectively (**Fig. 1H**). In addition, metabolome analysis showed that activated S/F-iTregs and nTregs consumed glucose and glutamine in the culture medium much less than activated Tconvs or C-iTregs (**Fig. 1I**). The result indicated less dependency of the activation/expansion of the former on these substances and correlated with lower expression of glucose transporter genes such as *Slc2a1* in S/F-iTregs and nTregs (**Fig. 1E**). Furthermore, S/F-iTregs, which suppressed both CD4^+^ and CD8^+^ responder T cells (**Fig. S1I**), were more potent than C-iTregs and equivalent to nTregs in *in vitro* suppressive activity (**Fig. 1J**).

Thus, S/F-iTregs similar to nTregs in Treg-related gene expression, epigenetic features and their potent suppressive function, hence stable in phenotype and function, can be efficiently produced from naive Tconvs.

### Generation of S/F-iTregs from effector/memory Tconvs

Next, we attempted to convert CD44^+^CD62L^low^ effector/memory CD4^+^ Tconvs from normal mouse lymph nodes into Foxp3^+^ Tregs using the protocol shown in Fig. 1 (**Fig. 2A-C**). While effector/memory CD4^+^ Tconvs were resistant to C-iTreg induction, either Senexin A addition or CD28 signal deprivation alone enhanced the generation of Foxp3^+^ cells from effector/memory CD4^+^ Tconvs; and combining both measures augmented the generation and also upregulated Foxp3, CD25 and CTLA-4 expression (**Fig. 2A-C** and **S2A**). Moreover, the secondary stimulation following the resting culture with IL-2 alone generated S/F-iTregs from effector/memory CD4^+^ Tconvs as efficiently as from naive Tconvs shown in Fig. 1. When S/F-iTregs induced from naïve Tconvs or effector/memory Tconvs (designated S/F-iTreg-TNs and S/F-iTreg-TEMs, respectively) were compared, the latter showed higher expression of Foxp3 and CTLA-4 (**Fig. 2D**). While the two S/F-iTreg populations were comparable in *in vitro* suppressive activity and in the expression levels of many activation-related markers (**Fig. 2E** and **S2B**), S/F-iTreg-TEMs showed relatively high in *Il2*, *Ikzf2*, *Il10*, *Il4*, and *Ifng* transcription (although TPM of *Il2* was less than 3) when compared with S/F-iTreg-TNs (**Fig. 2F**). In addition, RNA-seq and flow cytometric analysis revealed that S/F-iTreg-TNs were CD62L^high^, while S/F-iTreg-TEMs were Tim-3^high^ and TIGIT^high^ (**Fig. 2F** and **S2C**), indicating that S/F-iTregs maintained some of the immunological features present before conversion.

**Fig. 2.**
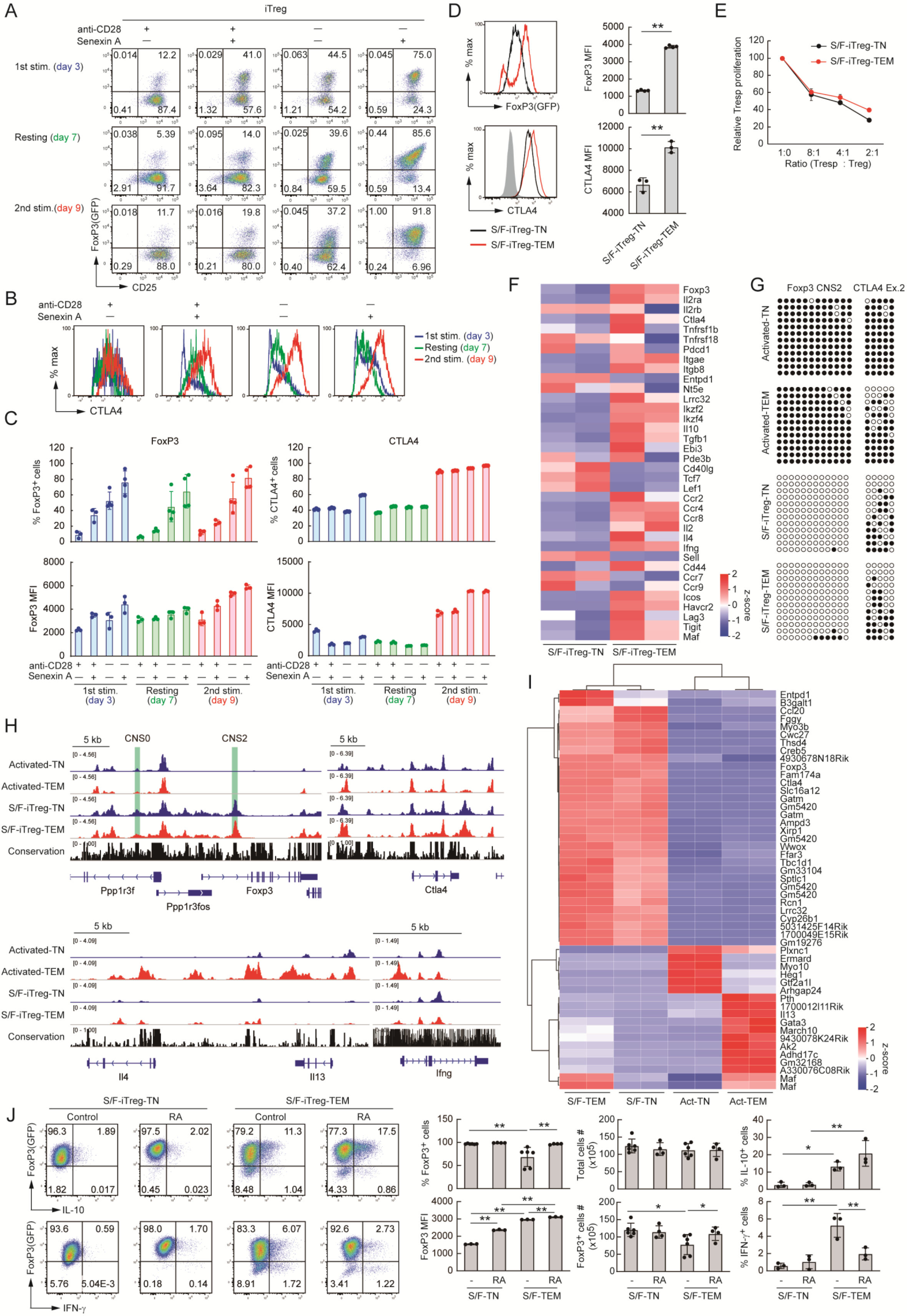
Generation of S/F-iTregs from effector/memory CD4^+^ Tconvs. (**A**) Protocol of iTreg induction from effector/memory (CD44^hi^CD62L^lo^) CD4^+^ Tconv suspensions from LNs of normal mice and Foxp3/CD25 staining of CD4^+^ Tconvs at each step of iTreg induction (representative of 3 independent experiments). (**B**) CTLA-4 staining of CD4^+^ Tconvs at each step of iTreg induction shown in (A). (**C**) Percentages and MFIs of Foxp3^+^ cells, CD25^+^ cells and CTLA-4^+^ cells assessed by flow cytometry at each step of iTreg induction (n=3-4). (**D**) Foxp3 and CTLA-4 expression of S/F-iTregs generated from naïve or effector CD4^+^ Tconvs (S/F-iTreg-TN and S/F-iTreg-TEM, respectively). MFI of FoxP3 and CTLA-4 assessed by flow cytometry (n=3-4). Unpaired t-test for statistical analysis (** p<0.01). (**E**) Suppressive activity of S/F-iTreg-TNs and S/F-iTreg-TEMs. Proliferation of Treg-cocultured CFSE-labelled responder CD4^+^ Tconvs (Tresp) in the presence of anti-CD3 and APCs was assessed and expressed as percentages compared with proliferation of Tresp cells alone as 100% (n=3). (**F**) Gene expression pattern of Treg-related molecules in S/F-iTreg-TNs and S/F-iTreg-TEMs (n=2). (**G**) DNA hypomethylation of *Foxp3* CNS2 and *Ctla4* Ex2 regions assessed by bisulfite sequencing. Filled circle; methylated CpG, unfilled circle; demethylated CpG. Representative of 2 or 3 independent experiments. (**H**, **I**) Chromatin accessibility assessed by ATAC-seq. Heat map shows top 50 differential accessible chromatin peaks (n=2). (**J**) Effects of retinoic acid (RA) in the second stimulation in (A). The numbers of total and FoxP3^+^ cells, and FoxP3, IL-10 and IFN-γ expression assessed by flow cytometry (n=3-6). Vertical bars denote mean±SD in C, D, and J. ANOVA followed by SNK test for statistical analysis (* p<0.05, ** p<0.01).

As for epigenetic status, both populations showed stable *Foxp3* CNS2 DNA hypomethylation (**Fig. 2G**) and high chromatin accessibility in the *Foxp3* and *Ctla4* gene loci (**Fig. 2H**). Notably, unlike activated Tconvs, especially activated effector/memory Tconvs, neither S/F-iTreg-TNs nor S/F-iTreg-TEMs showed high chromatin accessibility in the *Il4*/*Il13* and *Ifng* gene loci (**Fig. 2H**). The top 50 differential ATAC-seq peaks including the ones in these gene loci also indicated a distinct chromatin reprograming in S/F-iTregs compared with activated Tconvs (**Fig. 2I** and **S2D**). Further, both S/F-iTreg-TNs and S/F-iTreg-TEMs exhibited stable Foxp3 expression after *in vivo* transfer into non-lymphopenic mice (**Fig. S2E**).

Despite showing similar function and phenotype described above, S/F-iTreg-TEMs were less (∼60%) in cell yield than S/F-iTreg-TNs (**Fig. 2J**). To improve the yield of the former, we assessed the effects of RA, rapamycin, and butylate, which are all known to enhance Foxp3 expression (24–26), on S/F-iTreg-TEM generation (**Fig. S2F**), and found that RA indeed increased their yield (**Fig. 2J**). RA also promoted Foxp3 expression in both S/F-iTreg-TEMs and S/F-iTreg-TNs (**Fig. 2J**). Notably, a sizable (∼15%) fraction of the former produced IL-10, contrasting with less (∼2%) IL-10 production by the latter, which correlated with more active transcription of *Il10* and *Maf* by the former (**Fig. 2F**). Addition of RA to the secondary stimulation maintained the number of IL-10-producing cells, and also significantly reduced the number of IFN-γ-producing cells among S/F-iTreg-TEMs (**Fig. 2J** and **S2G**).

Collectively, the protocol for S/F-iTreg production from naive CD4^+^ T cells was also able to convert effector/memory CD4^+^ T cells into S/F-iTreg-TEMs, with improvement of the conversion by RA, which augmented their expansion and IL-10 production with reduction of IFN-γ production.

### Enhancement of S/-iTreg function by pharmacological reagents

In addition to RA, we searched for pharmacological reagents that could enhance and stabilize Treg function. Based on the previous findings on glucocorticoid-dependent expansion of nTregs and their high intracellular cAMP levels that contribute to Treg-mediated suppression (35, 36), we assessed dibutyryl-cAMP (dbcAMP), a cAMP analog, and dexamethasone, and found that they synergistically increased the percentage of Foxp3^+^IL-10^+^ iTregs (**Fig. S3A**). The effect was only observed when the reagents were added to the secondary stimulation for S/F-iTreg induction, and was not found with C-iTreg induction presumably because these compounds strongly inhibited T cell proliferation required for C-iTreg generation (**Fig. S3A**). In addition to IL-10 production, dbcAMP and dexamethasone augmented CTLA-4 expression by S/F-iTregs (**Fig. S3B**), and significantly inhibited their production of inflammatory cytokines especially TNF-α (**Fig. S3C**), albeit their *in vitro* suppressive activity was not significantly enhanced (**Fig. S3D**). RNA-seq analysis indeed revealed that dbcAMP/dexamethasone-treated S/F-iTregs enhanced the expression of Treg-signature genes including *Ctla4* and *Il10* (**Fig. S3E**). GO pathway analysis indicated that differentially expressed genes (DEGs) between dbcAMP/dexamethasone-treated and non-treated S/F-iTregs were attributed to down-regulation of the genes involved in glycolysis and other metabolic processes (e.g., *Slc2a1*, *Hk2* and *Gapdh*) and up-regulation of the ones engaged in immune responses or developmental processes (e.g., *Satb1*, *Pdcd1* and *Ctla4*) (**Fig. S3F** and **S3G**).

Ascorbate (Vitamin C), a co-enzyme for Tet demethylating enzymes, is known to enhance *Foxp3* CNS2 CpG hypomethylation (37–39). Both ascorbate-treated and non-treated S/F-iTregs showed a similar level of *Foxp3* CNS2 hypomethylation (**Fig. S3H**). Notably, Bcl2 expression was higher in ascorbate-treated S/F-iTregs (**Fig. S3I**), indicating that ascorbate might improve the survival of S/F-iTregs after *in vivo* transfer.

Taken together, S/F-iTregs and nTregs are similar in metabolic conditions and sensitivity to pharmacological reagents (such as RA, dbcAMP, dexamethasone, and ascorbate) that can control Treg-type metabolisms and epigenomic patterns to induce and augment nTreg-like functions in S/F-iTregs.

### S/F-iTreg induction from differentiated effector Th cells

We next attempted to convert Th1, Th2, and Th17 effector cells into Tregs by the protocol described above, with addition of RA to the secondary stimulation. For Th1 conversion, CD4^+^ Tconvs from draining lymph nodes of DNFB-induced contact hypersensitivity lesions were enriched for CD44^+^CXCR3^+^ cells, which were T-bet^high^, and subjected to the conversion (**Fig. 3A-F**). The generated Foxp3^+^ cells, designated S/F-iTreg-Th1s, were as CD25^high^ and complete in *Foxp3* CNS2 DNA hypomethylation as S/F-iTreg-TNs generated from naive CD44^low^CXCR3^-^ cells (**Fig. 3A**). The conversion efficiency to yield S/F-iTreg-Th1s was slightly (∼20%) less compared with the generation of S/F-iTreg-TNs. The former were higher than the latter in CTLA-4 expression, but slightly less suppressive *in vitro* suppression assay. Notably, ∼10% of S/F-iTreg-Th1s expressed IFN-γ compared with <1% of S/F-iTreg-TN cells; ∼20% of the former expressed T-bet compared with <1% of the latter, with ∼20% of both populations expressing CXCR3.

**Fig. 3.**
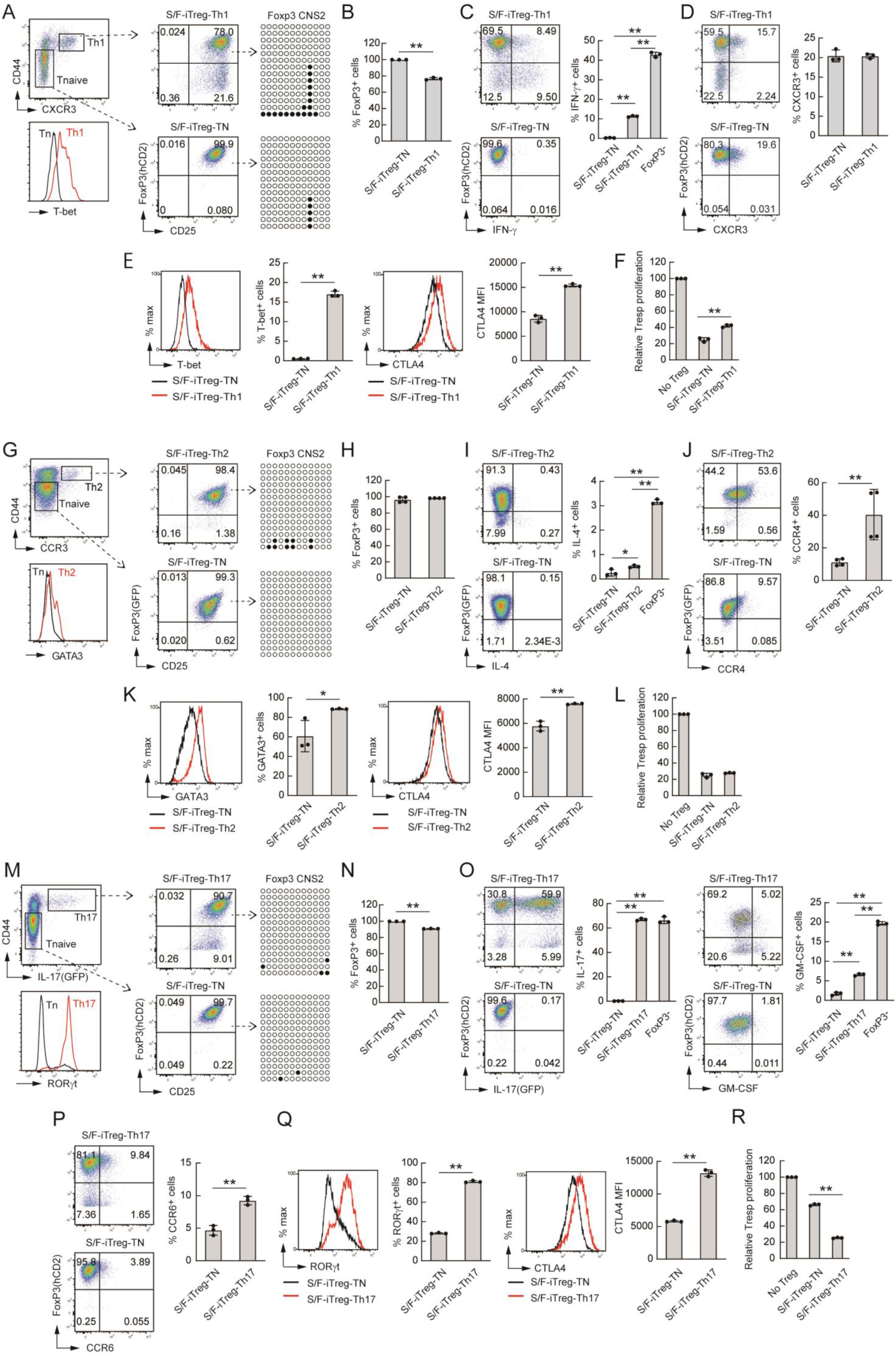
S/F-iTreg induction from differentiated effector Th cells. (**A-E**) S/F-iTreg induction from CXCR3^+^ Tconvs in draining LNs of epicutaneously DNFB-immunized mice. Percentages and MFIs of Foxp3^+^, IFN-γ^+^, CXCR3^+^, T-bet^+^ and CTLA-4^+^ cells assessed by flow cytometry (n=3). DNA hypomethylation of *Foxp3* CNS2 region assessed by bisulfite sequencing. Representative of 3 independent experiments. (**F**) Suppressive activity of iTregs. Proliferation of Treg-cocultured CFSE-labelled responder CD4^+^ Tconvs (Tresp) in the presence of anti-CD3 and APCs was assessed and expressed as percentages compared with proliferation of Tresp cells alone as 100% (n=3). (**G-K**) S/F-iTregs induction from CCR3^+^ T cell suspensions from spleen of mice immunized i.p. with OVA in alum adjuvant. Percentages and MFIs of Foxp3^+^, IL-4^+^, CCR4^+^, GATA3^+^ and CTLA-4^+^ cells assessed by flow cytometry (n=3-4). DNA hypomethylation of *Foxp3* CNS2 region assessed by bisulfite sequencing. Representative of 3 independent experiments. (**L**) Suppressive activity of iTregs as shown in (F) (n=3). (**M-Q**) S/F-iTreg induction from IL-17^+^ T cell suspensions from draining LNs of IL-17/GFP reporter mice immunized s.c. with MOG in CFA. Percentages and MFIs of Foxp3^+^, IL-17^+^, GM-CSF^+^, CCR6^+^, ROR-γt^+^ and CTLA-4^+^ cells assessed by flow cytometry (n=3). DNA hypomethylation of *Foxp3* CNS2 region assessed by bisulfite sequencing. Representative of 3 independent experiments. (**R**) Suppressive activity of iTregs as shown in (F) (n=3). Vertical bars denote mean±SD in B-F, H-L, and N-R. Unpaired t-test for statistical analysis in B, D-F, H, J-L, N and P-R (** p<0.01). ANOVA followed by SNK test for statistical analysis in C, I and O (* p<0.05, ** p<0.01). Filled circle; methylated CpG, unfilled circle; demethylated CpG in A, G and M.

For Th2 conversion, splenic CD4^+^ T cells from BALB/c mice intraperitoneally immunized with OVA in aluminum hydroxide were enriched for CCR3^+^ cells, which were largely GATA3^+^ (**Fig. 3G-L**). Such S/F-iTreg-Th2s could be generated as efficiently as S/F-iTreg-TNs, with complete *Foxp3* CNS2 demethylation, high CTLA-4 expression, and an equivalent *in vitro* suppressive activity. The former were also higher than the latter in the expression of GATA3 (∼90% vs ∼60%), IL-4 (∼0.5% vs ∼0.2%), and CCR4 (∼40% vs ∼10%).

Similarly, S/F-iTreg-Th17 cells with complete *Foxp3* CNS2 demethylation and high CTLA-4 expression were generated from GFP^+^CD4^+^ T cells in the draining lymph nodes of IL-17-GFP reporter mice immunized with MOG in CFA for inducing experimental allergic encephalomyelitis (**Fig. 3M-R**). They were higher in CTLA-4 expression and more potent in suppressive activity than S/F-iTreg-TNs. Notably, S/F-iTreg-Th17s were higher than S/F-iTreg-TNs in RORγt (∼80% vs ∼30%) and CCR6 (∼10% vs ∼5%) expression; and some (∼60%) were still producing IL-17 at an equivalent level as non-converted Foxp3^-^CD4^+^ T cells, although their production of GM-CSF was much lower than the latter (∼7% vs ∼20%).

In these experiments, the procedures (i.e., the use of the chemokine receptors or reporter GFP) for purifying each Th population before Treg conversion are different. Nevertheless, the results demonstrate that pathogenic effector Th cells can be converted into S/F-iTregs, which retained some of the phenotype of the original Th populations.

### Induction of antigen-specific S/F-iTregs and their specific *in vivo* activation by antigen stimulation

To prepare antigen-specific S/F-iTregs and assess their function, we generated S/F-iTregs from OVA-specific DO11.10 (DO) TCR transgenic CD4^+^ T cells (which can be detected by the KJ1.26 TCR clonotype-specific mAb, ref 40) as in Fig. 2; i.e., DO T cells were OVA stimulated in the presence of CTLA4-Ig to block CD80/CD86 co-stimulatory molecules on APCs, while control DO C-iTregs similarly produced by OVA stimulation without CTLA4-Ig and without Senexin A (**Fig. 4A**). DO S/F-iTregs were higher than C-iTregs in the ratios of FoxP3 and CTLA-4 expressing cells, the expression levels of these molecules and the degree of *Foxp3* CNS2 hypomethylation (**Fig. 4A** and **4B**). They potently suppressed DO Tconv proliferation upon OVA or anti-CD3 stimulation, while WT S/F-iTregs exerted suppression only upon anti-CD3 polyclonal stimulation (**Fig. 4C**). To determine then whether antigenic stimulation could preferentially generate antigen-specific S/F-iTregs, DO CD4^+^ Tconvs and WT Tconvs were mixed at 1:9 ratio and subjected to S/F-iTreg generation with OVA or anti-CD3 stimulation (**Fig. 4D**). OVA stimulation generated and expanded KJ1.26^+^ DO S/F-iTregs to assume ∼80% of Foxp3^+^ cells with some conversion of KJ1.26^-^ Tconvs as well, while anti-CD3 stimulation induced DO and WT S/F-iTregs at the original ratio of 1:9. *In vivo*, almost all DO S/F-iTregs prepared from Thy-1.2 DO Foxp3-reporter mice showed stable Foxp3 expression after transfer into syngeneic Thy-1.1 congenic mice and subsequent OVA immunization, whereas the majority (>80%) of DO C-iTregs had lost Foxp3 expression, consequently proliferating better than DO S/F-iTregs (**Fig. 4E**). When DO and WT S/F-iTregs were compared for their expansion/survival and phenotype after cell transfer and OVA immunization, DO S/F-iTregs expanded and survived better and were higher in Foxp3 and CTLA-4 expression at 2 weeks post immunization; they were still detectable as Foxp3^+^CTLA-4^+^ cells 4 weeks later when WT S/F-iTregs were scarcely detected, although the expression levels of these molecules gradually reduced in DO S/F- iTregs without further antigenic stimulation (**Fig. 4F** and **4G**). In addition, when DO or WT S/F-iTregs were transferred to DO RAG^-/-^ mice and OVA immunized, only the former showed potent suppressive activity as hindrance of the activation of the host DO T cells to become CD44^+^ and as suppression of their proliferation as Ki-67^+^ cells (**Fig. 4H** and **4I**). In contrast to such DO S/F-iTregs, few nTregs prepared from DO mice and OVA stimulated *in vitro* were detected 2 weeks after cell transfer and OVA immunization, suggesting that they might have lower *in vivo* proliferative activity, higher susceptibility to apoptosis, or both, upon antigenic stimulation (**Fig. S4A**). Such DO S/F-iTregs after *in vivo* OVA stimulation more closely resembled activated DO nTregs in transcriptome (**Fig. S4B**). In addition, when S/F-iTreg-Th1s and S/F-iTreg-TNs (**Fig. 3A-3F**) generated from CXCR3^+^ or CXCR3^-^ CD4^+^ T cells in OVA-immunized DO TCR transgenic mice were transferred to BALB/c mice and subsequently OVA immunized, both populations stably expressed Foxp3 and scarcely produced IFN-γ when examined 2 weeks after transfer (**Fig. S4C**).

**Fig. 4.**
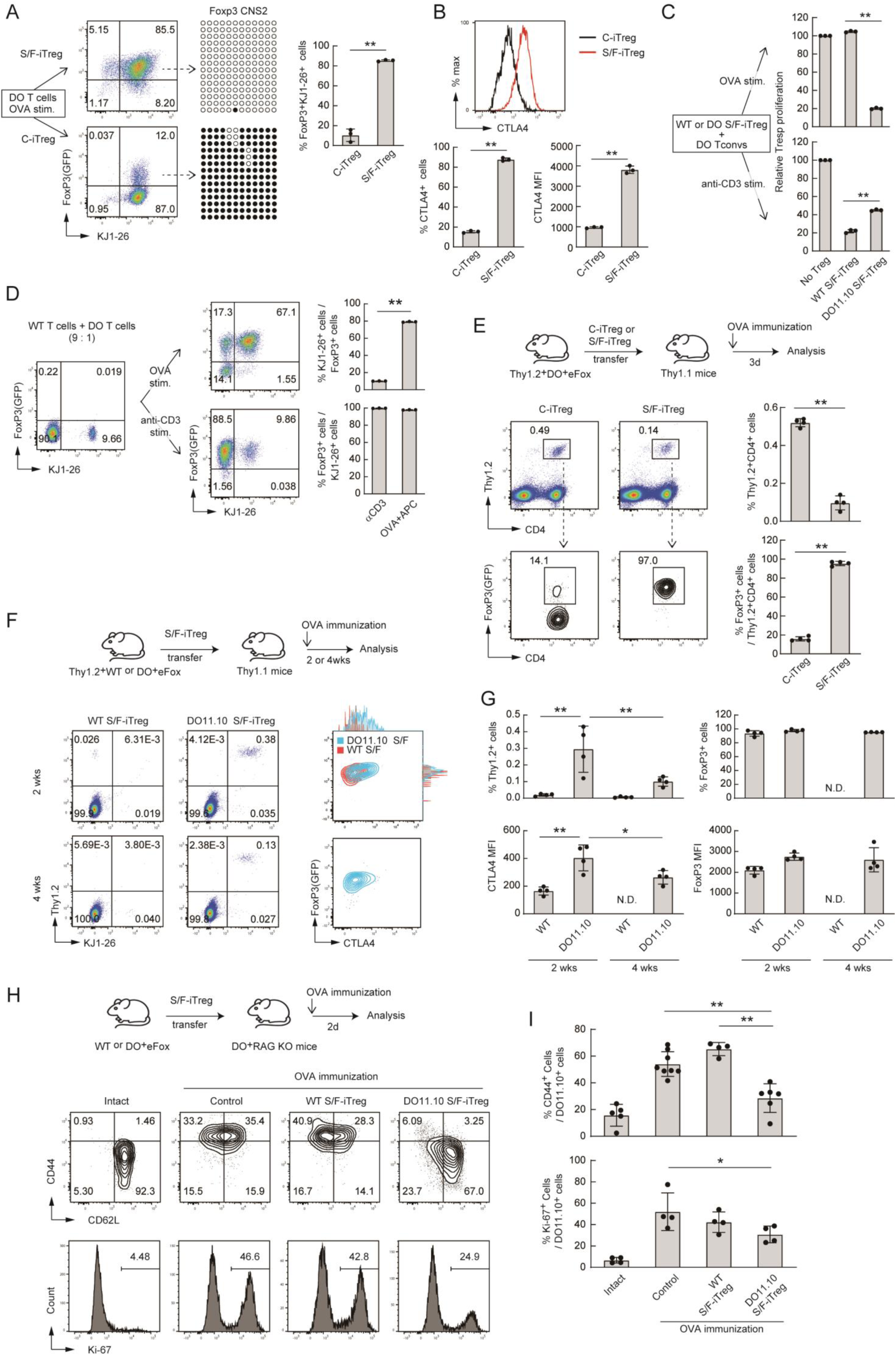
Induction of antigen-specific S/F-iTregs and their specific activation by *in vivo* antigenic stimulation. (**A**) iTreg induction from CD4^+^ T cell suspensions from LNs of DO11.10 TCR-transgenic mice. (**B**) Percentages and MFIs of Foxp3^+^ cells, KJ1-26^+^ cells, and CTLA-4^+^ cells were assessed by flow cytometry (n=3) and DNA hypomethylation of Foxp3 CNS2 region by bisulfite sequencing. Filled circle; methylated CpG, unfilled circle; demethylated CpG. (**C**) Suppressive activity of Treg populations. Proliferation of Treg-cocultured CFSE-labelled responder CD4^+^DO11.10^+^ T cells (Tresp) in the presence of anti-CD3 and APCs or OVA and APC was assessed and expressed as percentages compared with proliferation of Tresp cells alone as 100% (n=3). (**D**) S/F-iTregs induction from mixed suspensions of CD4^+^ T cells from WT- and DO11.10 mice. WT and DO CD4^+^ T cells at 9:1 ratio were converted to S/F-iTregs by OVA peptide or polyclonal anti-CD3 mAb stimulation. Percentages of Foxp3^+^ cells and KJ1-26^+^ cells were assessed by flow cytometry (n=3). (**E**) *In vivo* stability of OVA-specific iTregs. Thy1.2/DO11.10 Tregs were transferred into Thy1.1 WT mice and immunized with OVA. Percentages of Foxp3^+^ and Thy1.2^+^ cells in dLNs were assessed by flow cytometry 3 days later (n=4). (**F** and **G**) *In vivo* persistency of WT and DO11.10 S/F- iTregs. Thy1.2 Tregs were transferred into Thy1.1 WT mice and immunized with OVA. Percentages and MFI of Foxp3^+^ cells, Thy1.2^+^ cells and CTLA-4^+^ cells in dLNs were assessed by flow cytometry 2 or 4 weeks later (n=4). (**H** and **I**) *In vivo* suppressive activity of WT and DO11.10 S/F-iTregs (2×10^5^) assessed by adoptive transfer into DO11.10/Rag2 KO mice followed by OVA immunization. Percentages of CD62L^+^, CD44^+^ and Ki-67^+^ cells in the Foxp3^-^ fraction assessed by flow cytometry 2 days after immunization (n=4-8). Vertical bars denote mean±SD in A-E, G and I. Unpaired t-test for statistical analysis in A-E (** p<0.01). ANOVA followed by SNK test for statistical significance for G and I (* p<0.05, ** p<0.01).

These results indicate that antigen-specific Tconvs in the CD4^+^ Tconv compartment can be efficiently converted *in vitro* into S/F-iTregs by the S/F-iTreg generating procedure involving specific antigen stimulation. Upon antigen stimulation *in vivo* after cell transfer, such antigen-specific S/F-iTregs acquired Treg-type transcriptome much closer to that of antigen-specific nTregs when compared with S/F-iTregs before transfer (**Fig. 1F**), can expand specifically, survive long without producing inflammatory cytokines, and exert potent antigen-specific suppression.

### Suppression of murine inflammatory bowel disease (IBD) by S/F-iTregs

IL-10 secretion is required for Tregs to maintain mucosal tissue homeostasis (41, 42). However, S/F-iTreg-TNs produced less IL-10 compared to S/F-iTreg-TEMs (**Fig. 2J**). We therefore assessed dbcAMP/dexamethasone-treated, hence IL-10 producing S/F- iTregs to suppress IBD in a mouse model in which Thy1.2 Tregs were co-transferred with Thy1.1 naïve (i.e., CD45RB^high^) CD4^+^ Tconvs into RAG2^-/-^ mice (**Fig. 5A**). Both control and dbcAMP/dexamethasone-treated S/F-iTreg-TNs, as well as nTregs, prevented clinical and histological disease development, whereas C-iTregs did not (**Fig. 5A, 5B** and **S5A**). S/F-iTregs and nTregs maintained high percentages of Foxp3^+^ cells among transferred Tregs even 6 weeks after transfer, whereas C-iTregs had lost their Foxp3 expression by then (**Fig. 5C** and **5D**). Transcriptome analysis at the time revealed that S/F-iTregs retained a Treg-specific gene expression pattern as seen in nTregs (**Fig. S5B**), and exhibited potent suppressive activity *in vitro* (**Fig. S5C**). It was also noted that the transferred S/F-iTregs and nTregs enhanced the development of Foxp3^+^ cells from co-transferred CD45RB^high^ CD4^+^ Tconvs, which were Foxp3^-^ (**Fig. 5C** and **5D**). Unlike C-iTregs, the transferred S/F-iTregs and nTregs themselves scarcely produced IFN-γ and effectively suppressed IFN-γ production by co-transferred Tconvs (**Fig. 5E** and **5F**). Moreover, transferred Tregs, especially dbcAMP/dexamethasone-treated S/F-iTregs, elicited IL-10 production in co-transferred Tconvs (**Fig. 5G** and **5H**).

**Fig. 5.**
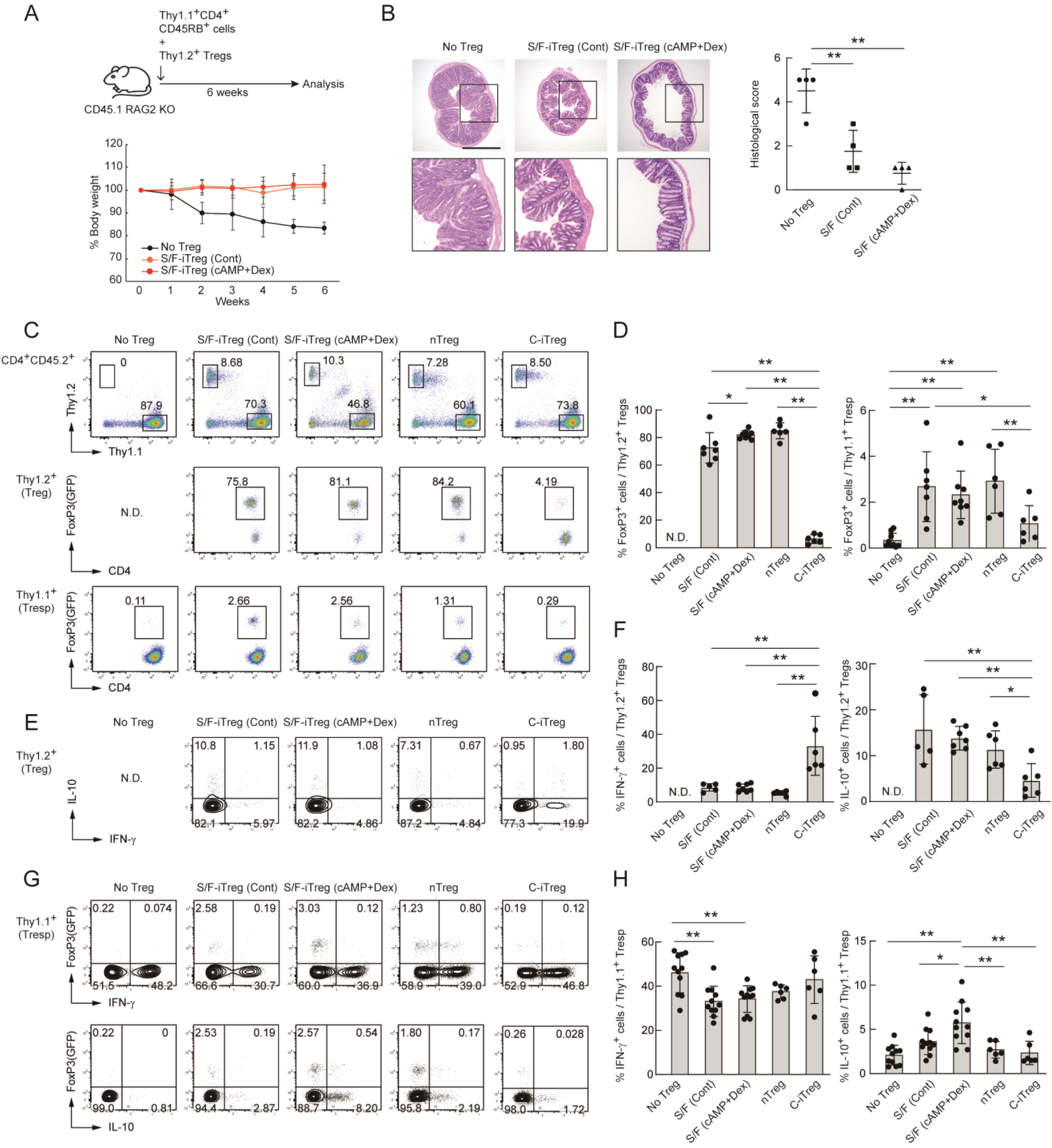
Suppression of murine inflammatory bowel disease by IL-10-secreting S/F-iTregs. (**A**) Colitis development in CD45.1/RAG2 KO mice by transfer of Thy1.1^+^CD4^+^CD45RB^hi^ naïve T cells with or without Thy1.2^+^ Tregs. (**A**) Relative body weight expressed as percentages of the weight before cell transfer (n=6-8). (**B**) Histological analysis of large intestine at the end point (n=4). Bar = 1 mm. (**C**) Representative staining of Thy1.2^+^ or Thy1.1^+^ cells in recipient mice for Foxp3 expression. (**D**) Percentages of FoxP3^+^ cells among Thy1.1 naïve T cells and Thy1.2 infused Tregs in mLNs assessed by flow cytometry (n=6-9). (**E**, **F**) IFN-γ^+^ and IL-10^+^ cells among Thy1.2^+^ infused Tregs. Representative staining (E) and percentages of cytokine-positive cells (F) (n=5-7). (**G, H**) IFN-γ^+^ and IL-10^+^ cells among Thy1.1^+^ naïve T cells in mLNs. Representative staining (G) and percentages of cytokine-positive cells (H) (n=6-11). Vertical bars denote mean±SD in B, D, F and H. ANOVA followed by Turkey’s post hoc test for statistical analysis in B (** p<0.01). ANOVA followed by SNK test for statistical analysis in D, F and H (* p<0.05, ** p<0.01).

Thus, S/F-iTregs, especially those producing IL-10, are able to suppress IBD directly and also, in part, by *de novo* inducing pTregs or eliciting IL-10 production from Tconvs apparently as an “infectious tolerance” (43).

### Treatment of murine graft-versus-host disease (GvHD) by S/F-iTregs

For induction of GvHD, BALB/c (H-2^d^) bone marrow cells and splenocytes were co-transferred into X-irradiated C57BL/6 (B6) (H-2^b^) hosts, which succumbed to severe GvHD with the mortality of 80% within 2 weeks post transfer. Co-transfer of S/F-iTreg-TNs prolonged the survival of the recipient mice in a dose-dependent manner while S/F- iTreg-TEMs were less effective (**Fig. 6A** and **6B**). For example, 1×10^6^ S/F-iTreg-TNs and nTregs equivalently suppressed GvHD, with ∼70% of the mice remaining alive over 30 days post transfer, while S/F-iTreg-TEMs were effective only at a large dose (2.5×10^6^ cells). The effect of S/F-iTregs to prolong survival was not affected by administration of the calcineurin inhibitor cyclosporine A, a current standard way for GvHD prophylaxis (**Fig. 6C**).

**Fig. 6.**
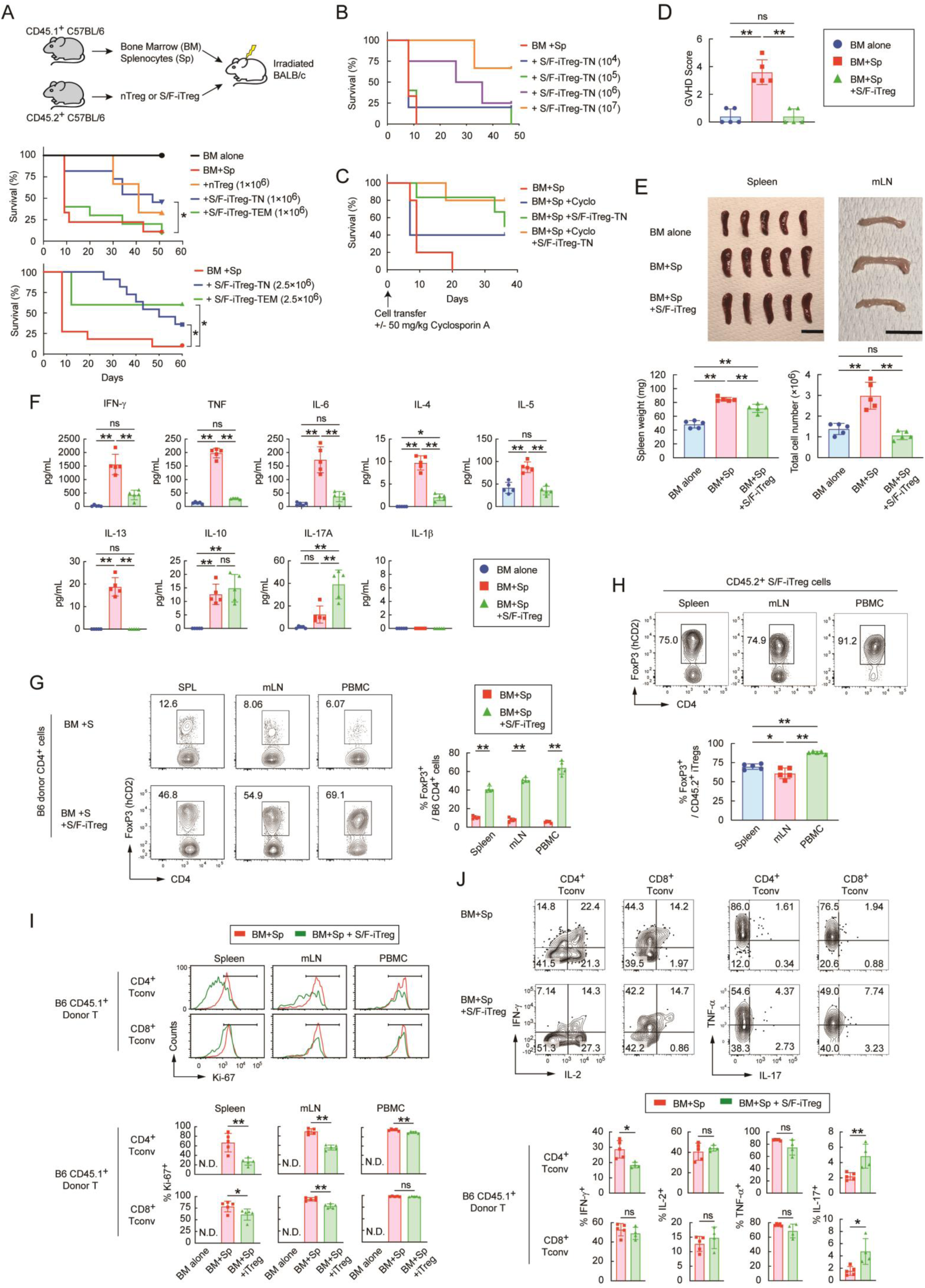
Suppression of allogenic GvHD in mice by S/F-iTregs prepared from naïve Tconvs. (**A**) Effects of adoptive Treg transfer on allogenic GvHD. 5×10^6^ BM cells and/or 2.5×10^6^ splenocytes (Sp) from CD45.1^+^Thy1.2^+^B6 mice were transferred into x-irradiated CD45.2^+^Thy1.1^+^BALB/c mice, together with or without indicated numbers of S/F-iTreg-TNs, S/F-iTreg-TEMs, or nTregs prepared from CD45.2^+^Thy1.2^+^B6 mice on day 0 (n=3-11 per group). (**B**) Dose effects of S/F-iTreg-TNs on GvHD. Indicated numbers of S/F- iTregs were co-transferred with BM+Sp on day 0 (n=3-5). (**C**) Effects of S/F-iTregs and cyclosporin A on GvHD prophylaxis. Cyclosporine A (CsA) (50 mg/kg) was administered intraperitoneally on day 0 with or without 2.5×10^6^ S/F-iTreg-TNs (n=5). (**D**) Pathology of GvHD 5 days after cell transfer (n=5-6). (**E**) Representative photograph of spleen (left panel) and ss (right panel) 5 days post transfer. The weight of spleens and total cell number of mLNs are shown (n=5). (**F**) Serum cytokine profiles 5 days after cell transfer. Serum samples collected from inferior vena cava were measured with Cytometry Beads Array (CBA) (n=4-5). (**G**) Treg increase after S/F-iTreg transfer. Percentages of Foxp3^+^ cells among B6-derived total CD4^+^ T cells in the spleen (SPL), mLNs, and peripheral blood (PB), assessed by flow cytometry 5 days after transfer (n=4-5). (**H**) Stability of transferred Tregs. Percentages of Foxp3^+^ cells among CD45.2^+^CD4^+^B6 cells assessed by flow cytometry (n=4-5). (**I**) Suppressive activity of S/F-iTregs. Percentages of Ki-67^+^ cells in CD8^+^ cells or CD4^+^ Tconvs collected from spleen, mLNs, or peripheral blood assessed by flow cytometry 5 days after transfer (n=5). (**J**) Inflammatory cytokine secretion from donor Sp. Percentages of cytokine positive cells in CD4^+^Foxp3^-^CD45.1^+^ or CD8^+^CD45.1^+^ donor B6 cells in mLNs were assessed by flow cytometry 5 days after transfer (n=4-5). Vertical bars denote mean±SD in D-J. Gehan-Breslow-Wilcoxon test in A-C, unpaired t-test in G and I or ANOVA followed by Turkey’s post hoc test in D-F and H for statistical analysis (* p<0.05, ** p<0.01). Bar = 10 mm.

At the cellular level, S/F-iTreg-TNs, which significantly suppressed GvHD when assessed 5 days after cell transfer (**Fig. 6D**), prevented splenomegaly and lympho-adenopathy, hindering the increase in the cellularity of the lymphoid organs (**Fig. 6E**). They also suppressed the elevation of inflammatory cytokines (e.g., IFN-γ, TNF-α, IL-6, IL-4, IL-5, and IL-13) while IL-17A was up-regulated, IL-10 being hardly changed, and IL-1β undetectable (**Fig. 6F**). The suppressed cytokine profile persisted on days 15 and 28 **(Fig. S6A)**. Notably, the IL-17A elevation on day 5 was transient and subsequently declined to near the detection limit by day 15. Histologically, S/F-iTreg-TN transfer alleviated lymphocytes infiltration around the portal area of the liver (data not shown).

The transfer of 2.5×10^6^ S/F-iTreg-TNs increased tenfold the percentage of Foxp3^+^ cells among CD4^+^ T cells in the peripheral blood, and 3∼6-fold in the spleen and mLNs (**Fig. 6G**). More than 80% of the transferred S/F-iTreg-TNs maintained Foxp3 expression in the circulation, and more than 60% in the spleen and mLNs (**Fig. 6H**). The expression of Ki-67 in donor-derived CD4^+^ and CD8^+^ T cells was significantly reduced primarily in the spleen and mLNs, to a lesser extent in the peripheral blood, suggesting that S/F-iTregs exerted immune suppression mainly in the secondary lymphoid tissues (**Fig. 6I**). Moreover, transferred S/F-iTregs significantly hampered the development of IFN-γ^+^CD4^+^ Th1 cells, with an increase in Th17 cells, in mLNs, correlating with the serum cytokine pattern (see above) (**Fig. 6J**). *In vitro* suppression assay in the presence of inflammatory cytokines revealed robust suppressive function of S/F-iTreg-TNs even in the Th1 and/or Th17 cytokine milieu (i.e., in the presence of IL-12, IL-6, and TGF-β). However, while S/F-iTreg cells effectively prevented Th1 differentiation, they failed to reduce the ratio of Th17 cells among responder Tconvs **(Fig. S6B)**.

Taken together, S/F-iTregs, especially S/F-iTreg-TNs, are able to suppress GvHD effectively by preferentially limiting Th1 over Th17 differentiation.

### Generation of human S/F-iTregs

We next attempted to generate S/F-iTregs from CD4^+^CD25^-^ Tconvs in human peripheral blood by the use of a similar protocol for murine S/F-iTreg generation, with addition of ascorbate (**Fig. 7A**). Compared with murine S/F-iTregs, the conversion rate of human Tconvs to FOXP3^+^ cells was more variable (average ∼80%) (**Fig. 7B**); however, modifications of the culture medium by adding RA and reducing glucose and glutamine concentrations by half increased the rate to ∼90%, with ∼25-fold increase in the number of FOXP3^+^ cells in 9 days from the start of the culture (**Fig. 7C**). These treatments also upregulated FOXP3 and CTLA-4 expression, producing FOXP3^+^CTLA-4^+^ cells as ∼80% of the yielded cells (**7D**, **Fig. S7A** and **S7B**). S/F-iTregs thus generated showed a gene expression profile similar to that of activated nTregs (**Fig. 7E**). They also exhibited Treg type epigenetic modifications such as CpG hypomethylation of the FOXP3 CNS2 region at a comparable level as nTregs (**Fig. 7F**). Ascorbate further enhanced the hypomethylation, which progressed mainly during the second stimulation (**Fig. S7C**). Chromatin accessibility was similar between S/F-iTregs and nTregs at the FOXP3 CNS2 region (**Fig. 7G**). *In vitro* suppressive activity of S/F-iTregs was significantly higher than C-iTregs and nTregs (**Fig. 7H**). In addition, S/F-iTregs were stable in FOXP3 expression and scarcely produced proinflammatory cytokines when restimulated *in vitro* in the presence of various proinflammatory cytokines (**Fig. 7I** and **S7D**). The TCR repertoire assessed by the frequencies of T cells expressing particular TCR Vβ subfamilies was not altered before and after S/F-iTreg conversion (**Fig. S7E**).

**Fig. 7.**
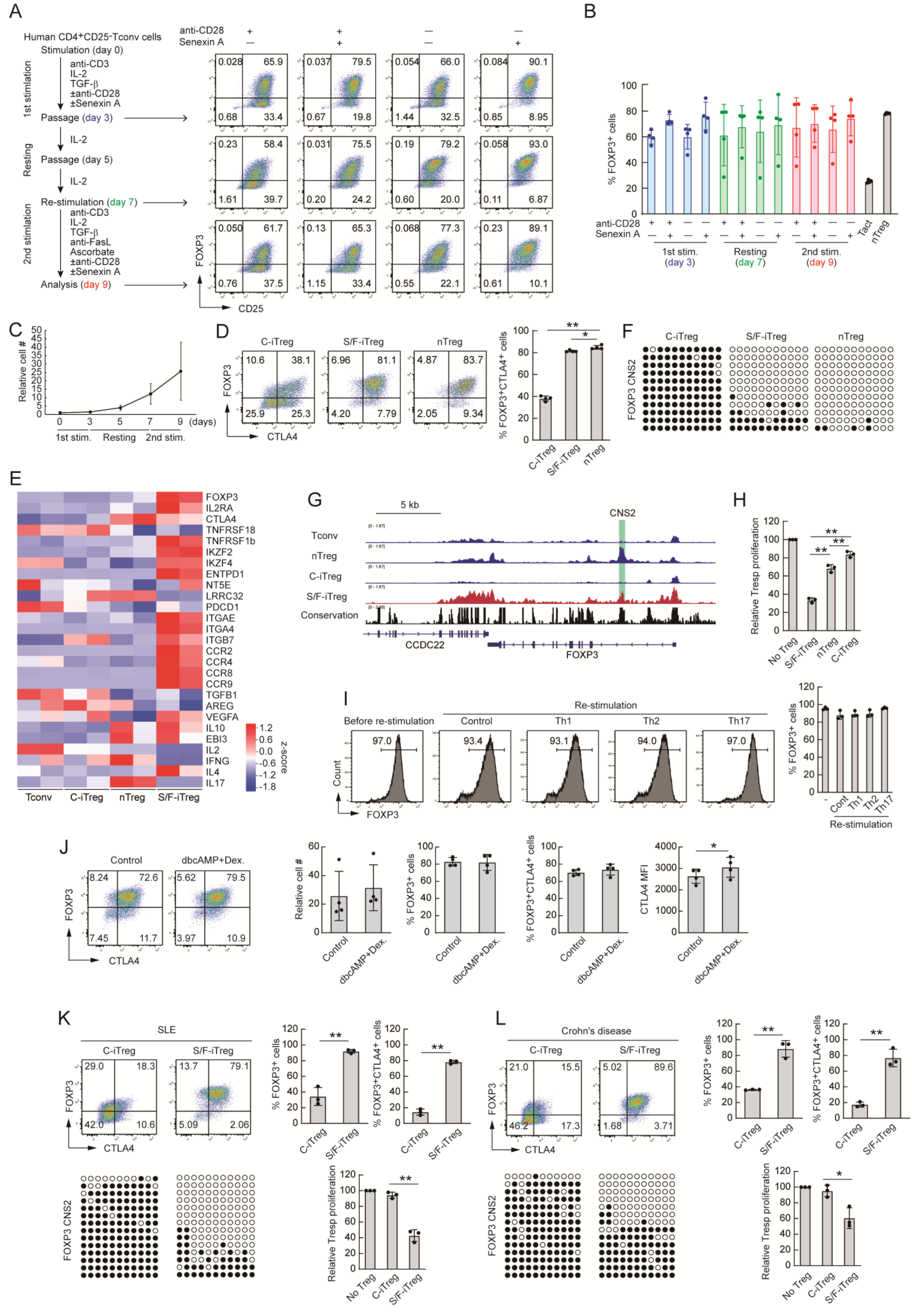
Generation of human S/F-iTregs. (**A**) Protocol for iTreg cell induction from human CD4^+^CD25^-^ Tconv suspensions prepared from PBMCs of healthy donors and FOXP3/CD25 staining of CD4^+^ T cells at each step of Treg cell induction. (**B**) Percentages of FOXP3^+^ cells at each step of iTreg induction (n=4). (**C**) S/F-iTreg cell yield relative to the initial cell number of Tconvs for Treg conversion (n=4). (**D**) Percentage of FOXP3^+^ and CTLA4^+^ cells in S/F-iTregs, conventional iTregs (C-iTregs), and nTregs assessed by flow cytometry (n=4). (**E**) Gene expression pattern of Treg-related molecules in *in vitro* activated Tconvs, C-iTregs, *in vitro* stimulated nTreg cells, and S/F-iTregs (n=2). (**F**) DNA hypomethylation of *FOXP3* CNS2 region assessed by bisulfite sequencing. Representative of 3 independent experiments. (**G**) Chromatin accessibility assessed by ATAC-seq. Heat map shows top 50 differential accessible chromatin peaks. Representative of 2 independent experiments. (**H**) Suppressive activity of human S/F-iTregs as shown in Fig. 1J for mouse S/F-iTregs (n=3). (**I**) *In vitro* stability of S/F-iTregs assessed by anti-CD3 mAb stimulation in the presence of inflammatory cytokines. Percentage of FOXP3^+^ cells assessed by flow cytometry (n=3). (**J**) The effect of dbcAMP (20 μM) and Dexamethasone (10 nM) on human S/F-iTregs. Cell numbers relative to the initial cell number, percentages and MFI of FOXP3^+^ cells and CTLA-4^+^ cells assessed by flow cytometry (n=4). (**K**) S/F-iTregs and C-Tregs induced from CD4^+^ Tconvs of a male SLE patient. DNA hypomethylation of *FOXP3* CNS2 region and percentage of FOXP3^+^ and CTLA-4^+^ cells assessed by bisulfite sequencing and flow cytometry, respectively. Suppressive activity was assessed as shown in (G) (n=3). (**L**) S/F-iTregs and C-Tregs induced from CD4^+^ Tconvs of a female Crohn’s disease patient. DNA hypomethylation of *FOXP3* CNS2 region (note: the maximum demethylation rate is ∼50% because this is female patient sample) and percentage of FOXP3^+^ and CTLA-4^+^ cells were assessed as shown in (K). Suppressive activity was assessed as shown in (G) (n=3). Vertical bars denote mean±SD in B, C, D and H-L. ANOVA followed by SNK test for statistical analysis in D and H (* p<0.05, ** p<0.01). Paired t-test for statistical analysis in J, and unpaired t-test for statistical analysis in K and L (* p<0.05, ** p<0.01). Filled circle; methylated CpG, unfilled circle; demethylated CpG in F, K and L.

As shown with murine S/F-iTregs, addition of dbcAMP and dexamethasone further enhanced FOXP3 and CTLA-4 expression in human S/F-iTregs without inhibiting cell proliferation (**Fig. 7J**). The treatment also improved the generation of CTLA-4^+^ S/F- iTregs from effector/memory Tconvs, with higher IL-10 production (**Fig. S7F-S7H**). These modifications of S/F-iTreg production indeed enabled generating S/F-iTregs from the peripheral blood of patients with autoimmune/inflammatory diseases, for example, systemic lupus erythematosus (SLE) (**Fig. 7K**), Crohn’s disease (**Fig. 7L**), and pemphigus vulgaris (44). CCR6^+^Th17-like cells, which constituted ∼1/3 of CD4^+^ T cells in PBMCs of SLE patients (45–47), could also be converted into S/F-iTregs (**Fig. S7I**); such effector-derived S/F-iTregs showed a higher reactivity to autologous APCs (**Fig. S7J**).

Thus, S/F-iTregs functionally and phenotypically similar to nTregs can be efficiently generated from autoimmune/inflammatory disease patients as well as healthy individuals even if the patients are under immunosuppressive treatment with steroid.

## Discussion

We have shown in this study that antigen-primed effector/memory as well as naïve Tconvs can be converted into antigen-specific Tregs by providing two conditions. One is the antigenic stimulation of Tconvs in the presence of CDK8/19 inhibitor, TGF-β, and IL-2 to induce strong Foxp3 expression, the other being the deprivation of CD28 co-stimulation during iTreg generation to confer Treg-type epigenetic changes (30, 31). Repeating this Treg conversion process with intermittent resting cultures with IL-2 alone, efficiently yielded S/F-iTregs even from cytokine-producing Th cells. The results demonstrate that iTregs functionally similar to nTregs can be generated by manipulating specific signaling pathways, including the TGF-β/TGF-β receptor/SMAD, the IL-2/IL-2 receptor/STAT5, and the CDK8/19/STAT5 pathways for Foxp3 induction, and inhibition of the CD28/PKC/NFκB pathway for Treg-type DNA hypomethylation. The efficacy of S/F-iTreg conversion can be further increased by creating the cell culture conditions more favorable for Tregs than for Tconvs in cell expansion/survival because of distinct metabolic features of the former (e.g., less metabolic dependency of Tregs on glycolysis and glutaminolysis, hence enrichment of Tregs by reducing glucose and glutamine concentrations in the culture medium) (48). The efficacy can also be enhanced by adding RA, dbcAMP, and dexamethasone to enrich Foxp3^+^ cells, augment their suppressive activity, induce more IL-10 production, and reduce inflammatory cytokine production. In addition, depriving CD28 signal and supplementing ascorbate can synergistically enhance *Foxp3* CNS2 DNA hypomethylation, ensuring stable Foxp3 expression. By incorporating these procedures into a single protocol, we have shown that mouse and human CD4^+^ Tconvs, whether in naïve or activated effector/memory states, can be converted into S/F-iTregs with high (more than 90%) efficiency.

For iTregs to be as stably suppressive as nTregs, they need as minimum requirements to express CD25 and CTLA-4 highly and constitutively and not to secrete IL-2, as supported by the following findings. Treg-specific deletion of CD25 in Tregs or systemic IL-2 neutralization severely impairs Treg function and survival, causing severe autoimmune/inflammatory diseases, while provision of IL-2 neutralizes Treg-mediated suppression *in vitro* (16, 17, 49, 50). Treg-specific CTLA-4 deficiency abrogates Treg suppression and evokes fatal autoimmunity (6). Indeed, a combination of IL-2 non-production, high constitutive expression of CTLA-4, and TCR stimulation (which up-regulates the expression of CD25, hence the high affinity IL-2 receptor, as well as LFA-1 and other adhesion molecules) can confer Treg-like *in vivo* and *in vitro* suppressive activity on Tconvs without Foxp3 expression (51). S/F-iTregs in mice and humans show this molecular expression/non-expression pattern through acquiring Treg-specific transcriptional and epigenetic changes. In addition, Treg-type epigenome alterations, especially DNA demethylation, are found exclusively in Foxp3^+^ cells in S/F-iTreg generation despite the fact that Foxp3 expression and Treg-specific DNA hypomethylation can be separately induced in Tconvs subjected to Treg conversion (13, 31). Further understanding of how the two molecular events occur simultaneously to generate S/F-iTregs and for that matter nTregs may enable more robust generation of the former and reveal the molecular basis of the function and development of the latter.

S/F-iTregs can be generated from mouse and human effector/memory fraction Tconvs and also from mouse cytokine-secreting Th1, Th2, and Th17 cells, while conserving, to various extents, the expression of the respective Th-specific TFs, cytokines, and chemokine receptors. They contrasted with S/F-iTreg-TNs generated from the same mice as the latter scarcely produced relevant signature cytokines or chemokine receptors. Further, unlike activated effector/memory Tconvs, S/F-iTreg-TEMs did not show high chromatin accessibility at the *Il4*/*Il13* and *Ifng* gene loci. Functionally, IFN-γ secreted from S/F-iTreg-Th1 cells slightly impaired *in vitro* suppression, whereas IL-17 from S/F-iTreg-Th17 cells did not. In addition, S/F-iTreg-TEMs tended to produce more IL-10 than S/F-iTreg-TNs. Addition of RA, dbcAMP, and dexamethasone further increased IL-10 production and significantly reduced IFN-γ-production in S/F-iTreg-TEMs, along with augmented CTLA-4 expression. Moreover, repetition of the S/F-iTreg inducing cell culture further reduced IFN-γ and IL-17 production in S/F-iTregs (our unpublished data). On the other hand, the conserved expression of Th-specific TFs such as Tbet, GATA3, and RORγt in respective S/F-iTreg-Ths may sustain the expression of Th-specific chemokine receptors such as CXCR3, CCR3/CCR4, and CCR6, thereby enabling the respective S/F-iTreg-Th cells to migrate to the sites affected by relevant inflammation types. Thus, advantages and disadvantages of retaining the original Th-specific immunological properties after S/F-iTreg conversion need to be further investigated in order to enable S/F-iTregs to effectively suppress immune responses depending on the site and type of inflammation.

We have shown that S/F-iTregs specific for a particular antigen are more potent than polyclonal S/F-iTregs in suppressing Tconvs reactive with the antigen upon antigenic stimulation. Further supporting the finding, self-antigen-specific S/F-iTregs prepared from CD4^+^ T cells specific for desmograin-3, the skin antigen solely targeted in the skin autoimmune disease pemphigus vulgaris, specifically expanded in the skin-draining lymph nodes and effectively suppressed disease progression in a mouse model of pemphigus vulgaris (44). These results underline the critical importance of antigen specificity of Tregs in immune suppression, raising the issue of how much of polyclonal S/F-iTregs prepared from an autoimmune disease patient possess relevant self-antigen specificities. There is accumulating evidence that normal individuals harbor in the peripheral blood potentially hazardous Tconvs recognizing self-antigens targeted in autoimmune diseases (52–54). It is apparently the frequency (i.e., clonal expansion) and the immunological state (i.e., naïve, anergic, or activated) of such potentially pathogenic T cells that are different between healthy individuals and autoimmune disease patients (53, 54). It is therefore a key advantage of S/F-iTregs that, with possible clonal expansion of pathogenic self-reactive Tconvs in the peripheral blood of a patient, polyclonal S/F- iTreg conversion of the patient’s CD4^+^ Tconvs, especially the activated effector/memory fraction, results in enrichment of the corresponding self-antigen-specific S/F-iTregs. The ratio of antigen-specific Tconvs present before conversion is indeed maintained after polyclonal conversion to S/F-iTregs. In many autoimmune/inflammatory diseases, however, the target antigen is unknown or multiple, the main pathogenic target specificity being obscure, and the predominant pathogenic specificities different among patients (depending in part on HLA haplotypes). Nevertheless, polyclonal Treg conversion of the whole effector/memory T-cell fraction, including pathogenic Tconvs, can generate S/F- iTregs enriched for those possessing relevant disease-specific suppressive activity. Such patient-derived polyclonal S/F-iTregs would eventually achieve antigen-specific or disease-specific immune suppression because those S/F-iTregs stimulated by known or unknown self-antigens targeted in an autoimmune disease may show better *in vivo* expansion and longer survival than non-stimulated ones after adoptive cell transfer.

In contrast with the conceivable high efficacy of effector/memory-derived S/F-iTregs in antigen-specific immune suppression as discussed above, it is of note that S/F-iTregs from naïve Tconvs are more potent than those from effector/memory Tconvs in suppressing GvHD. This finding may reflect the fact that naïve Tconvs are more potent than effector/memory-type Tconvs in causing GvHD presumably because the former possess a broader and stronger alloreactivity than the latter (55–57). It further supports the notion that S/F-iTregs generated from disease-mediating Tconvs are more effective in disease suppression compared with those from non-mediating ones. These results collectively indicate that the generation of therapeutically effective antigen- or disease-specific S/F-iTregs from naïve or effector/memory Tconvs depends on the roles of Tconvs in mediating particular physiological and pathological immune responses.

In addition to these immunological characteristics of S/F-iTregs, they possess some distinct immunological properties when compared with other types of Foxp3^+^ Tregs. For example, nTregs can be synthetically bestowed with a particular antigen- or a disease-specificity by expressing engineered TCRs or chimeric antigen receptors (CARs) (reviewed in 58, 59). Both S/F-iTregs and engineered nTregs require antigen stimulation for their expansion and exertion of suppressive function; on the other hand, they need to avoid functional exhaustion due to chronic TCR stimulation. It has been shown mainly with CAR-T cells, and similarly with CAR-Tregs, that chronic tonic stimulation through CAR renders them dysfunctional (60, 61). In contrast, S/F-iTregs maintain proliferative activity and suppressive function despite continuous *in vitro* TCR stimulation and subsequent *in vivo* antigenic stimulation. One of the possible differences between S/F-iTregs and CAR-Tregs can be an attenuation of TCR-proximal signaling upon TCR stimulation in the former but not in the latter in which CAR-mediated strong signal may exceed physiological TCR signal. That is, upon TCR stimulation, S/F-iTregs as well as nTregs down-regulate the expression of Lck and ZAP-70 in a Foxp3-dependent manner, consequently attenuating the intensity of TCR signaling and thereby rendering them resistant to functional exhaustion to be incurred by chronic strong TCR stimulation (62, 63). In addition, such activated S/F-iTregs appear to be more resistant to activation-induced cell death than activated nTregs as suggested by higher expression of Bcl-2 family anti-apoptotic genes and lower expression of pro-apoptotic genes in the former. Furthermore, S/F-iTregs are apparently able to induce “infectious tolerance” (43) by converting Treg-suppressed Tconvs into Foxp3^+^ cells and IL-10-producing cells as effectively as nTregs at least under certain conditions (e.g., in a colitis model). It has been shown that activated Tregs down-regulate CD80/CD86 expression by APCs (6, 32, 33) and that deprivation of the CD80/CD86 costimulation facilitates the generation of functionally stable iTreg cells (31). It is therefore plausible that one-time introduction of S/F-iTregs may elicit consecutive *de novo* Treg induction to establish long-term suppression and tolerance. These immunological properties of S/F-iTregs may enable them to sustain stable dominant suppression over pathogenic Tconvs.

In conclusion, immunological and pharmacological manipulation of cell signaling pathways, without ectopic gene expression, is able to induce Treg-type gene transcription and epigenome formation in effector/memory as well as naive Tconvs. This technology can indeed generate functionally stable antigen-specific iTregs from Tconvs mediating physiological and pathological immune responses. S/F-iTregs thus prepared resemble nTregs immunologically in antigen specificity, survivability, suppressive function, functional stability in cytokine-rich inflammatory environments, cell migration to inflammation sites, and adaptability to inflammation types. Furthermore, sufficient numbers of S/F-iTregs can be obtained within a short period, which would allow their application in acute autoimmune conditions and organ transplant rejections. Thus, adoptive cell therapy with antigen-specific or disease-specific S/F-iTregs would be instrumental in treating autoimmune and other inflammatory diseases and preventing graft rejection in organ transplantation.

## Materials and Methods

### Study design

The aim of this study was to generate functionally stable iTregs from antigen-specific pathogenic effector/memory T cells for treating immunological diseases. They were produced in vitro by antigenic stimulation in the presence of cytokines (IL-2 and TGF-β) without co-stimulation and by repeating this process with intermittent resting cultute with IL-2 alone. The phenotype of such iTregs was assessed by flow cytometry, bisulfite sequence, RNA-seq, ATAC-seq, ChIP-seq and *in vitro* suppression assay. ATAC-seq and RNA-seq analysis was also performed to investigate the differences between iTregs generated from naive or effector/memory T cells. Bisulfite sequence, RNA-seq, ATAC-seq, ChIP-seq and metabolome analysis were performed on biological replicates.

Antigen-specific induction of iTregs was conducted using DO11.10 TCR transgenic T cells with OVA stimulation. Mouse models of IBD and GvHD were used to examine *in vivo* therapeutic potential of iTregs. Sufficient numbers (more than four) of adult mice with matched age (6 to 12 weeks) were used for each *in vivo* experiment. *In vitro* iTreg inductions and *in vivo* therapeutic effects were reproduced in two independent laboratories.

### Mice

C57BL/6, BALB/c mice were purchased from SLC or CLEA. DO.11.10 TCR transgenic mice, Rag2 KO mice, Thy1.1 congenic mice, Foxp3-eGFP (eFox) reporter mice and Foxp3-hCD2 reporter mice were previously described (22, 30, 31). IL-17-eGFP reporter mice were purchased from the Jackson Laboratory (strain #:018472). All procedures were performed in accordance with the National Institutes of Health Guide for the Care and Use of Laboratory Animals and approved by the Committee on Animal Research of Osaka University and Kyoto University.

### Antibodies and reagents

Antibodies are listed in Supplemental Table S1. Senexin A and dibutyryl-cAMP were purchased from Tocris Bioscience; rapamycin from Cell Signaling Technology; all trans-retinoic acid, butyrate, ascorbate and dexamethasone from Sigma; OVA (323–339) and MOG (35–55) peptides from MBL; and CTLA4-Ig (Abatacept; Orencia) from Ono Pharm co. Recombinant mouse IL-4, IL-6 and IL-12 were purchased from PeproTech. Cell Stimulation Cocktail (plus protein transport inhibitors) was purchased from Invitrogen for intracellular cytokine staining.

### Cell sorting and Flow cytometry analysis

Cell staining by fluorescence-conjugated antibodies and flow cytometry analysis was performed as previously described (31). To prepare cells for culture experiments, FACSAriaII (BD) was used for collecting particular populations. The definition of cell populations used are the following: naïve T cells, CD4^+^GFP^-^CD44^low^CD62L^high^; nTreg cells, CD4^+^GFP^+^; and effector/memory T cells, CD4^+^GFP^-^CD44^high^CD62L^low^. For intracellular staining, cell fixation and permeabilization were performed using the Foxp3/Transcription Factor Staining Buffer Set (Invitrogen) or BD Pharmingen Transcription Factor Buffer Set (BD) in accordance with the manufacturer’s instructions. For TCR repertoire analysis, cells were stained by Beta Mark TCR Vbeta repertoire kit (Beckman-Coulter). Samples were analyzed by FACSCantoII, FACSAriaII or FACSLyric frow cytometer.

### Cell culture, iTreg induction and resting culture

For mouse cell culture, we used RPMI1640 culture medium supplemented with 10% FCS (v/v), 60 µg/ml penicillin G, 100 µg/ml streptomycin and 0.1 mM 2-mercaptoethanol. For the induction of stable-iTregs, sorted 2×10^5^ CD4^+^ T cells were stimulated with plate-bound anti-CD3 mAb (coating at 10 μg/ml) in the presence of 50 U/ml of human IL-2 (Imunace 35; Shionogi Pharm Co.) and 5 ng/ml of human TGF-β1 (R&D), in 96 well flat bottom plate (Thermo Scientific, #167008). Soluble anti-CD28 mAb at 1 µg/ml was additionally used to generate conventional-iTregs. For resting culture, stimulated iTregs were collected, washed once, further cultured at 1×10^6^/ml cell concentration with fresh culture medium containing 100 U/ml of IL-2. Cells were split every 2 days and fresh IL-2-containing medium was added. For OVA peptide stimulation, 2×10^5^ DO11.10 CD4^+^ T cells were co-cultured with 2×10^4^ CD11c^+^ APCs in the presence of 5 μM of OVA peptide (MBL).

For S/F-iTregs induction, rested iTregs were re-stimulated with plate-bound anti-CD3 mAb (coating at 10 μg/ml) in the presence of 50 U/ml of human IL-2 (Imunace 35; Shionogi Pharm Co.), 5 ng/ml of human TGF-β1 (R&D) and anti-FasL mAb (5 μg/ml), in 96 well flat bottom plate (Thermo Scientific, #167008). Senexin A (Selleck, 5 μM), retinoic acid (AM80; TOCRIS, 1 μM), ascorbate (2-phospho-L-ascorbic acid, SIGMA, 10 μg/ml), dibutyryl cyclic AMP (Actosin; Alfresa Pharma Corp., 20 μM) and dexamethasone (Decadron; Sandoz Pharma K.K., 10 nM) were supplied as needed.

For human iTregs induction, we used RPMI1640 or D-Glucose and L-glutamine reduced RPMI generated by Shimadzu Diagnostics Corporation (Tokyo, Japan) supplemented with 10% FCS (v/v), 60 µg/ml penicillin G, 100 µg/ml streptomycin and 10 mM HEPES. CD4^+^CD25^-^ Tconv cells were sorted from PBMC purchased from Cellular Technology Limited, HemaCare Corporation and Astarte Biologics. Cells were cultured in 24 well flat bottom plate (Falcon, #351147), as described above for murine iTreg induction method, by using anti-human CD3, CD28 and FasL mAbs.

### Suppression Assay

CD4^+^ naïve T cells were labeled with violet proliferation dye (VPD; Cell Tracer Violet, Thermo Scientific) by the following protocol: cells were incubated at 1×10^6^ cells/ml with 5 μM reagents at room temperature for 5 min. Labeling reaction was quenched by adding 5 volumes of cold medium and incubated further for 20 min on ice. After washing cells once, 1×10^5^ labeled cells were co-cultured with 2×10^4^ antigen presenting cells and graded numbers of Tregs in the presence of 1 μg/mL anti-CD3 mAb or 5 μM OVA peptide. Percentages of cells dividing more than once were assessed by FACS after 72 hours.

### CpG methylation analysis by bisulfite sequencing

Bisulfite sequencing analysis was performed as previously described (31, 64). Cells were collected by FACSAriaII and DNA was extracted by phenol extraction followed by ethanol precipitation. When cells were fixed for intracellular staining, reverse-crosslinking reaction was performed overnight prior to gDNA extraction. The bisulfite conversion was carried out using the MethylEasy Xceed Rapid DNA Bisulphite Modification Kit (Human Genetic Signatures) by following manufacturer’s protocol. PCR primer sequences of commonly-methylated regions or Treg-specific demethylated regions are available in (13).

### RNA-sequencing and analysis

RNA-seq was performed as previously described (31). Cells were lysed in RLT buffer (Qiagen) containing 1% 2-Mercaptoethanol, followed by RNA reverse transcription by SMART-seq v4 Ultra Low Input RNA Kit for Sequencing (Clontech). Sequencing libraries were prepared using Kapa Library preparation kit for IonTorrent (KAPA) or KAPA HyperPlus Kits (KAPA) by following manufacturer’s protocol. Sequencing of cDNA libraries was performed on a IonS5 (Thermo Scientific) or DNBSEQ (MGI). TPM values were acquired using an RNA-seq integrative pipeline, ikra (v2.0.1), composed of Trim Galore! (v0.6.7), Salmon (v1.4.0), tximport (v1.6.0) with the reference Gencode M26 for mice and 37 for humans. Principal component analysis (PCA) and pathway analysis of DEGs were performed using iDEP (v0.96) (65).

### ATAC-sequencing analysis

ATAC-seq was performed as previously described (66). Briefly, sorted cells (up to 100,000 cells) were lysed using 100 µl of lysis buffer (0.01% digitonin, 0.1% NP-40, 0.1% Tween 20 in resuspension buffer; 10 mM Tris-HCl pH7.5, 100 mM NaCl, 3 mM MgCl2) for 3 min on ice. After discarding lysis buffer by centrifugation, Tn5 tagmentation was performed using Illumina Tagment DNA TDE1 Enzyme and Buffer Kits (Illumina) at 37°C for 30 min with shaking at 1,000 rpm. After purification using DNA Clean & Concentrator-5 (Zymo Research), tagmented DNA was amplified using NEBNext High-Fidelity PCR Master Mix (New England BioLabs). Prepared DNA libraries were cleaned up by DNA Clean & Concentrator-5 or Ampure XP (Beckman Coulter). Sequencing was performed using NextSeq500 or NovaSeq (Illumina).

For ATAC-seq analysis, sequenced reads were mapped to mm10 genome using Bowtie2 (v2.3.5) by following options; --very-sensitive. Mapped reads were sorted by Samtools (v1.6), then converted into bigwig files for visualization by bamCoverage (v3.1.3), included deepTools package with following options; -of bigwig --binSize 5 -p max --normalizeUsing CPM --smoothLength 15 –ignoreDuplicates. Peak calling was performed by MACS2 (v2.1.1.20160309) with following options; macs2 callpeak -t ${filename} -name ${outputfilename} -f BAM -g mm --SPMR --nolambda --nomodel - -shift -75 --extsize 150 --keep-dup all --bdg --call-summits. For further analysis, narrowPeak files were concatenated and sorted. Sorted narrowPeak file was merged using bedtools (v2.30.0) and then converted into SAF format file. SAF peak file was annotated using HOMER (v4.11). Count files were generated using featureCounts with following options; -a annotated.saf -F SAF -p -o counts.txt ${BAM}. Differential expression analysis was performed using DESeq2.

### ChIP-sequencing analysis

ChIP-seq experiments were performed as previously described (21, 22). Briefly, sorted cells were fixed using 1% Formaldehyde (Thermo Scientific) for 10 min for anti-Histone ChIP at room temperature. After nuclear extraction, chromatin lysate was fragmentated using Picoruptor (Diagenode), 30 sec sonication and 30 sec cooling for 7 cycles at 4°C, before immunoprecipitation. Immunoprecipitated chromatin lysate was reverse-crosslinked at 65°C for 24 h, followed by purification and library preparation using NGS library preparation kit for IonS5 (Thermo Scientific) or NEBNext Ultra II DNA Library Prep Kit for Illumina (Illumina) according to manufacturer’s instructions. Raw data were produced by IonS5 sequencer system (Thermo Scientific) or NextSeq500 (Illumina).

For ChIP-seq analysis, raw sequence reads were checked and trimmed using Trim- galore (v0.6.6) (Babraham Bioinformatics). Trimmed reads were mapped to mm10 genome using Bowtie2 (v2.3.5) by following options; --local --very-sensitive-local. Mapped reads were sorted by Samtools (v1.6), then converted into bigwig files for visualization by bamCoverage (v3.1.3), included deepTools package, with following options; -of bigwig --binSize 5 -p max --normalizeUsing CPM --smoothLength 15 – ignoreDuplicates. Peak calling was performed by MACS2 (v2.1.1.20160309) with following options; macs2 callpeak -t ${filename} -name ${outputfilename} -f BAM -g mm --SPMR --nolambda --nomodel --bdg --call-summits.

### DNFB immunization

For Th1 cell induction, mice were sensitized epicutaneously on day 0 by applying 100 μL of 0.5 % DNFB diluted in acetone on the abdominal skin. Draining lymph nodes were collected on day 2 to sort CD4^+^CD44^+^CXCR3^+^ cells as Th1-like cells.

### OVA immunization

For Th2 cell induction, mice were immunized intraperitoneally with 200 µg of OVA peptide in aluminium hydroxide gel (FUJIFILM Wako Chemicals.) on days 0 and 2. Spleen were collected on day 4 and sort CD4^+^CD44^+^CCR3^+^ cells as Th2-like cells.

For *in vivo* stability and functional assay of DO Tregs, Thy1.1 WT mice or DO11.10/RAG2 KO mice were immunized subcutaneously with 200 µg of OVA. Tregs (2×10^5^) were transferred just before immunization and draining lymph nodes were collected for flow cytometry analysis at various time points.

### MOG immunization

For Th17 cell induction, IL-17-GFP reporter mice were immunized subcutaneously with 100 μL of an emulsion containing 100 μg of MOG peptide in 50 μL of PBS and 50 μL of CFA on day 0. CFA was prepared by mixing incomplete Freund’s adjuvant (Difco) with 20 mg/ml of Mycobacterium tuberculosis H37RA (Difco). Pertussis toxin (List Biological Laboratories Inc., Campbell, CA) was injected on days 0 and 2 (100 ng/mouse, i.p.). Draining lymph nodes were collected on day 4 and sort CD4^+^CD44^+^GFP^+^ cells as Th17 cells.

### Colitis model

To induce inflammatory bowel disease, 2×10^5^ CD4^+^CD45RB^hi^ cells prepared from Thy1.1 mouse were injected into CD45.1/RAG2 KO mice. Same number of each Treg population was also transferred with CD45RB^high^ T cells at the same time. After 6 weeks transfer, histological analysis of colon tissue and flow cytometry analysis of mLNs were carried out. Histological scoring was measured by following criteria after the hematoxylin and eosin staining; no-disorder: 0, disorder of epithelium: 1, disorder of epithelium and partial ulcer: 2, disorder of epithelium and partial ulcer or partial cell invasion: 3, disorder of epithelium, partial ulcer, and partial cell invasion:4, severe colitis (disorder of epithelium, ulcer, and cell invasion whole colon tissue): 5.

### GvHD model

To induce GvHD in mice, donor-derived 5×10^6^ bone marrow cells and/or 2.5×10^6^ splenocytes from CD45.1/Thy1.2/H-2Kb C57BL/6 mice were intravenously injected into irradiated (5 Gly) CD45.2/Thy1.1/H-2Kd BALB/c recipients with or without CD45.2/Thy1.2/H-2Kb Tregs. For nTreg preparation, spleen and whole lymph nodes were harvested and total CD25^+^CD4^+^ T cells were isolated using CD4^+^CD25^+^ Regulatory T Cell Isolation Kit, mouse (Miltenyi Biotec), following the manufacturer instruction. When required, Cyclosporin A (50 mg/kg, FujiFilm Wako) suspended in saline was administered intraperitoneally to the recipients on day 0 of transfer. The survival and body weight change of recipients were recorded at least twice a week. GvHD pathology was evaluated based on the following criteria: diarrhea, skin redness, ruffled feathers, posture, and the amount of activity, with each criterion scored from 0 to 2. At the end point, whole blood was collected from central artery of the euthanized mice using 29G syringe. For serum isolation, the collected blood was loaded onto Bloodsepar (Immunuo-Biological Lab Co.), incubated at room temperature for 10 minutes and then centrifuged at 2400G, 4°C. The concentration of serum cytokines was determined using BD cytometry beads array (CBA assay). For mononuclear cell collection, the blood was loaded into 0.1% EDTA in PBS, followed by cell separation using Ficoll (Cytiva).

### Metabolome analysis

For metabolome analysis, supernatant was collected after iTreg induction or primary stimulation process. Relative amount of metabolites was quantified by LC/MS/MS Method Package for Cell Culture Profiling at Shimadzu Diagnostics Corporation (Tokyo, Japan).

### Statistics

Values were expressed as mean ± SD. Statistical significance was assessed by Paired or unpaired Student’s t-test (two groups), repeated-measures analysis of variance (ANOVA) followed by the Dunnett’s test (versus control), non-repeated-measures ANOVA followed by the Turkey’s post hoc test or Student-Newman-Keuls (SNK) test (multiple comparisons). A probability of less than 5% (P < 0.05) was considered statistically significant.

## Supplementary Materials

Fig. S1. Generation of S/F-iTregs from naïve CD4^+^ Tconvs.

Fig. S2. Generation of S/F-iTregs from naïve or effector/memory CD4^+^ Tconvs.

Fig. S3. Effects of cAMP analog, dexamethasone and ascorbate on S/F-iTreg function.

Fig. S4. *In vivo* expansion/survival of OVA-specific nTregs and S/F-iTregs upon OVA stimulation.

Fig. S5. *In vivo* suppressive activity of nTregs and C-iTregs on colitis.

Fig. S6. Suppressive function of S/F-iTregs under inflammatory cytokine production.

Fig. S7. Effects of retinoic acid, limitation of glucose and L-glutamine, cAMP analog and dexamethasone on human S/F-iTreg induction.

## Acknowledgments

We would like to thank Y. Nakamura, M. Hata, N. Okamoto, Y. Manabe, M. Matsuura and R. Ishii for technical assistance, C. Tay and Y.H. Oo for proof- reading of the manuscript and T. Shindo for technical guidance of GvHD model.

## Author contributions

NM, AS, RK and SS designed the project. NM, AS, RK and MA conducted the experiments and data analysis. NM, AS, RK and SS wrote the manuscript with contribution from all the authors.

## Data and materials availability

The raw data (fastq files) from the RNA-seq, ATAC-seq and ChIP-seq experiments have been deposited at the DDBJ Sequence Read Archive under accession no. PRJDB18080. All data needed to evaluate the conclusions in the paper are present in the paper or the Supplementary Materials.

