## supplemental Files for "Generation of antigen-specific and functionally stable Treg cells from effector/memory T cells for cell therapy of immunological diseases"

**The PDF file includes:**

Fig. S1. Generation of S/F-iTregs from naïve $CD4^+$ Tconvs.

Fig. S2. Generation of S/F-iTregs from naïve or effector/memory $CD4^+$ Tconvs.

Fig. S3. Effects of cAMP analog, dexamethasone and ascorbate on S/F-iTreg function.

Fig. S4. *In vivo* expansion/survival of OVA-specific nTregs and S/F-iTregs upon OVA stimulation.

Fig. S5. *In vivo* suppressive activity of nTregs and C-iTregs on colitis.

Fig. S6. Suppressive function of S/F-iTregs to control inflammatory cytokine production.

Fig. S7. Effects of retinoic acid, limitation of glucose and L-glutamine, cAMP analog and dexamethasone on human S/F-iTreg induction.

Table S1. List of antibodies.

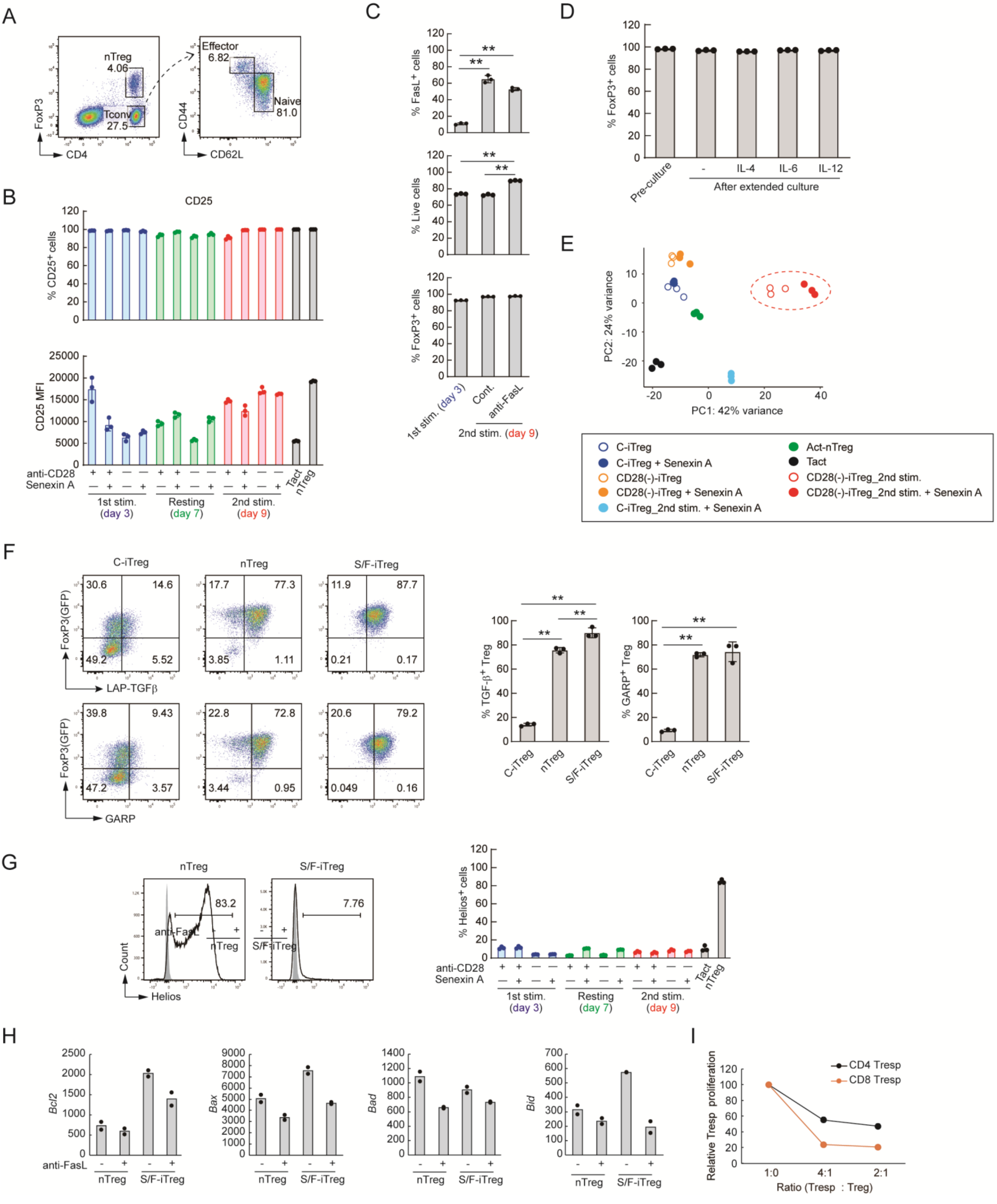
A
B
C
D
E
F
G
H
I
% FasL+ cells
% Live cells
% FoxP3+ cells
1st stim. (day 3)
Cont.
anti-FasL
2nd stim. (day 9)
Pre-culture
After extended culture
IL-4
IL-6
IL-12
CD25
% CD25+ cells
CD25 MFI
anti-CD28
Senexin A
1st stim.
(day 3)
Resting
(day 7)
2nd stim.
(day 9)
Tact
nTreg
PC1: 42% variance
PC2: 24% variance
C-iTreg
C-iTreg + Senexin A
CD28(-)-iTreg
CD28(-)-iTreg + Senexin A
C-iTreg_2nd stim. + Senexin A
Act-nTreg
Tact
CD28(-)-iTreg_2nd stim.
CD28(-)-iTreg_2nd stim. + Senexin A
C-iTreg
nTreg
S/F-iTreg
FoxP3(GFP)
LAP-TGFβ
GARP
% TGF-β+ Treg
% GARP+ Treg
Helios
Count
% Helios+ cells
Bcl2
Bax
Bad
Bid
anti-FasL
Relative Tresp proliferation
CD4 Tresp
CD8 Tresp
Ratio (Tresp : Treg)

**Fig. S1. Generation of S/F-iTregs from naïve $CD4^+$ Tconvs.**

(**A**) Gating strategy of FoxP3(eGFP)$^+$ cells, naïve ($CD62L^+CD44^{low}$), and effector ($CD62L^-CD44^{hi}$) $CD4^+$ T cells in lymph nodes of intact mice. (**B**) Percentages of $CD25^+$ cells assessed by flow cytometry at each step of Treg induction (n=3). (**C**) Percentage of live cells, $FoxP3^+$ cells and $FasL^+$ cells assessed by flow cytometry at day 3 and 9 (n=3). (**D**) *In vitro* stability of S/F-iTregs assessed by additional 3-day culture in the presence of inflammatory cytokines. Percentage of $FoxP3^+$ cells assessed by flow cytometry (n=3). (**E**) PCA of RNA-sequencing data shown in Fig. 1E, together with Senexin A-added samples (n=3). (**F**) Percentages of TGF-$\beta^+$ and $GARP^+$ cells assessed by flow cytometry (n=3). (**G**) Percentage of $Helios^+$ cells assessed by flow cytometry at each step of Treg induction (n=3). (**H**) Effects of anti-FasL mAb on apoptosis gene expression by S/F-iTregs or nTregs. Cells were stimulated with or without anti-FasL mAb *in vitro,* purified for live cells by FACS, and subjected to RNA-seq analysis to show TPM of the apoptosis-related genes shown (n=2). (**I**) In vitro suppressive function of S/F-iTregs on CD8 T cells. In vitro suppression assay was carried out by using APCs and $CD4^+$ or $CD8^+$ responder T cells (Tresp) in the presence of anti-CD3 stimulation (n=3). Vertical bars denote mean$\pm$SD. ANOVA followed by SNK test for statistical analysis in C and F (** $p<0.01$).

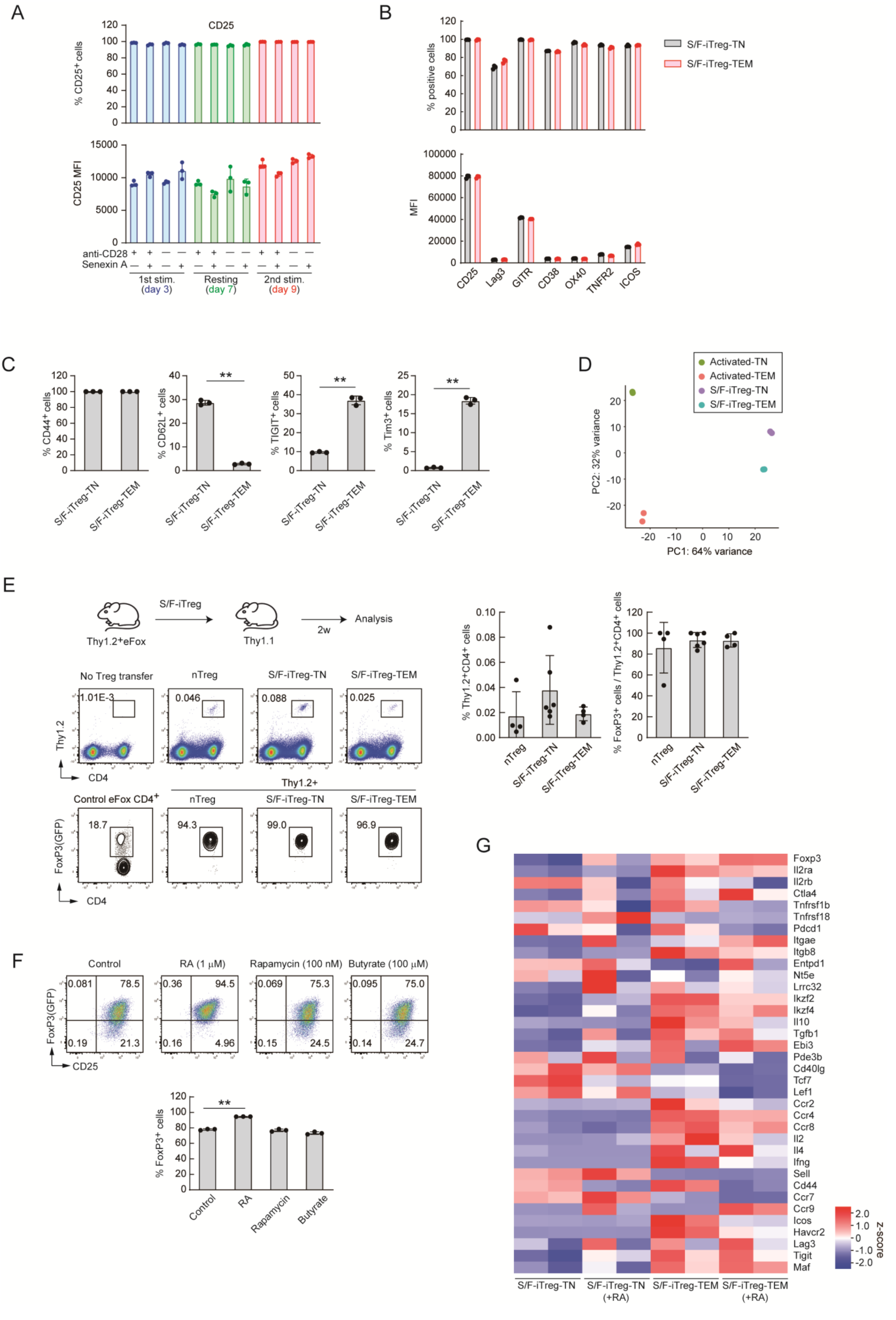
A
CD25
% CD25+ cells
CD25 MFI
anti-CD28
Senexin A
1st stim.
(day 3)
Resting
(day 7)
2nd stim.
(day 9)
B
% positive cells
MFI
S/F-iTreg-TN
S/F-iTreg-TEM
CD25
Lag3
GITR
CD38
OX40
TNFR2
ICOS
C
% CD44+ cells
% CD62L+ cells
% TIGIT+ cells
% Tim3+ cells
S/F-iTreg-TN
S/F-iTreg-TEM
D
Activated-TN
Activated-TEM
S/F-iTreg-TN
S/F-iTreg-TEM
PC2: 32% variance
PC1: 64% variance
E
S/F-iTreg
Thy1.2+eFox
Thy1.1
2w
Analysis
No Treg transfer
nTreg
S/F-iTreg-TN
S/F-iTreg-TEM
1.01E-3
0.046
0.088
0.025
Thy1.2
CD4
Control eFox CD4+
Thy1.2+
18.7
94.3
99.0
96.9
FoxP3(GFP)
% Thy1.2+CD4+ cells
% FoxP3+ cells / Thy1.2+CD4+ cells
F
Control
RA (1 μM)
Rapamycin (100 nM)
Butyrate (100 μM)
0.081
78.5
0.19
21.3
0.36
94.5
0.16
4.96
0.069
75.3
0.15
24.5
0.095
75.0
0.14
24.7
CD25
% FoxP3+ cells
RA
Rapamycin
Butyrate
G
Foxp3
Il2ra
Il2rb
Ctla4
Tnfrsf1b
Tnfrsf18
Pdcd1
Itgae
Itgb8
Entpd1
Nt5e
Lrrc32
Ikzf2
Ikzf4
Il10
Tgfb1
Ebi3
Pde3b
Cd40lg
Tcf7
Lef1
Ccr2
Ccr4
Ccr8
Il2
Il4
Ifng
Sell
Cd44
Ccr7
Ccr9
Icos
Havcr2
Lag3
Tigit
Maf
Z-score
S/F-iTreg-TN
S/F-iTreg-TN (+RA)
S/F-iTreg-TEM
S/F-iTreg-TEM (+RA)

**Fig. S2. Generation of S/F-iTregs from naïve or effector/memory $CD4^+$ Tconvs.**

(**A**) Percentages of $CD25^+$ cells assessed by flow cytometry at each step of Treg induction (n=3). (**B**) Percentage and MFI of surface markers in naïve- or effector/memory-derived S/F-iTregs assessed by flow cytometry (n=3). (**C**) Percentages of $CD44^+$, $CD62L^+$, Tim-$3^+$ and $TIGIT^+$ in naïve- or effector/memory-derived S/F-iTregs assessed by flow cytometry (n=3). (**D**) PCA of chromatin accessibility of activated naïve or effector/memory Tconvs and S/F-iTregs assessed by ATAC-seq (n=2). (**E**) *In vivo* stability of naïve- and effector/memory-derived S/F-iTregs. Thy1.2 Tregs were transferred into Thy1.1 WT mice. Percentages of $Foxp3^+$ cells and $Thy1.2^+$ cells in LNs were assessed by flow cytometry 2 weeks later (n=4-6). (**F**) Effects of known FoxP3 inducing compounds on effector/memory-derived S/F-iTregs. Percentage of $FoxP3^+$ cells assessed by flow cytometry (n=3). (**G**) Gene expression pattern of Treg-related molecules in S/F-iTreg-TNs and S/F-iTreg-TEMs with or without retinoic acid (n=2). Vertical bars denote mean$\pm$SD in A-C, E and F. Unpaired t-test for statistical analysis in C (** $p<0.01$). ANOVA followed by Dunnet's test for statistical analysis in F (** $p<0.01$ vs. control).

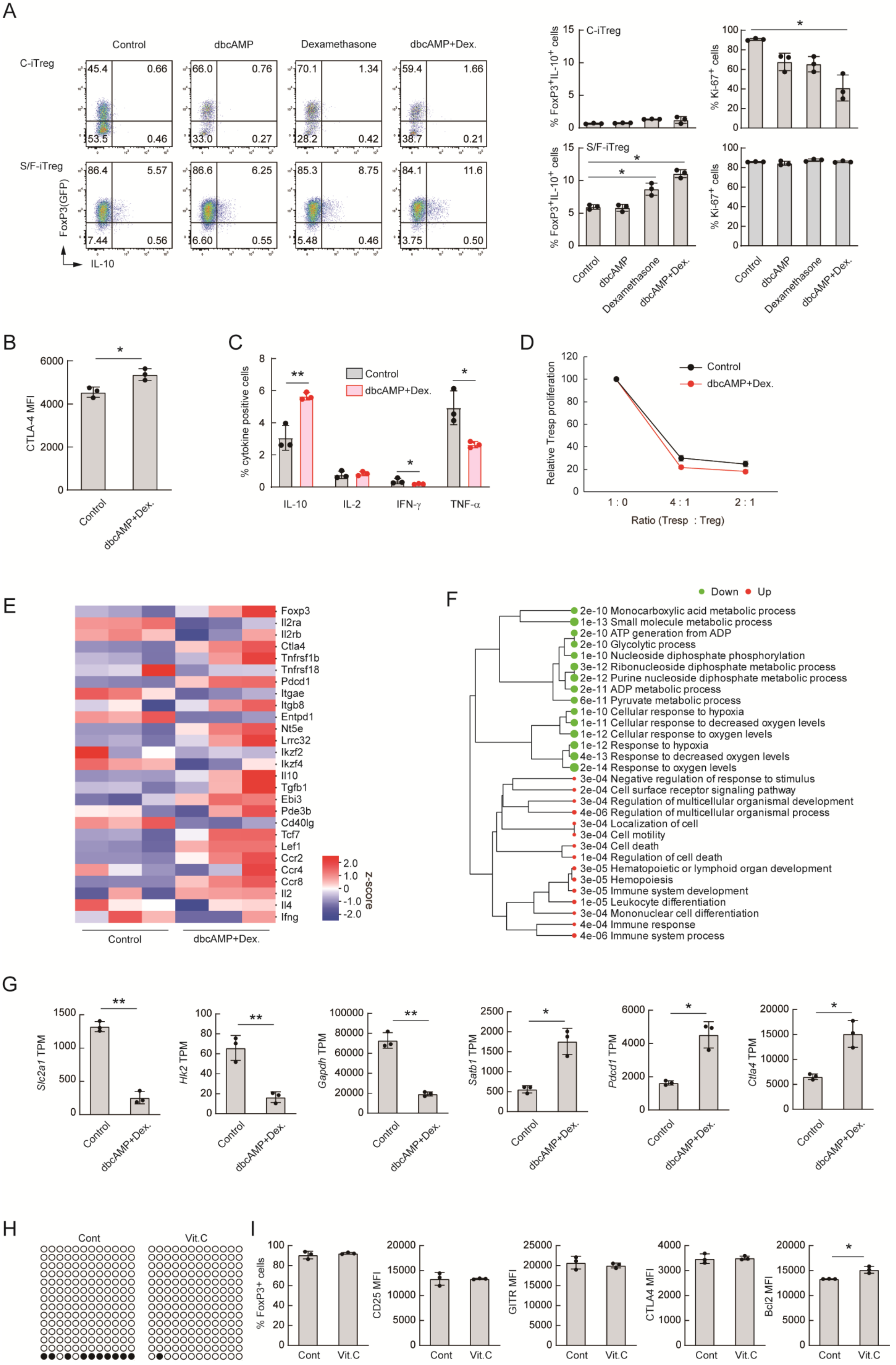
A
Control
dbcAMP
Dexamethasone
dbcAMP+Dex.
C-iTreg
45.4 0.66 53.5 0.46
66.0 0.76 33.0 0.27
70.1 1.34 28.2 0.42
59.4 1.66 38.7 0.21
S/F-iTreg
86.4 5.57 7.44 0.56
86.6 6.25 6.60 0.55
85.3 8.75 5.48 0.46
84.1 11.6 3.75 0.50
FoxP3(GFP)
IL-10
% FoxP3+IL-10+ cells
% Ki-67+ cells
C-iTreg
S/F-iTreg
Control
dbcAMP
Dexamethasone
dbcAMP+Dex.
B
CTLA-4 MFI
Control
dbcAMP+Dex.
C
% cytokine positive cells
Control
dbcAMP+Dex.
IL-10
IL-2
IFN-γ
TNF-α
D
Relative Tresp proliferation
Control
dbcAMP+Dex.
1 : 0
4 : 1
2 : 1
Ratio (Tresp : Treg)
E
Foxp3
Il2ra
Il2rb
Ctla4
Tnfrsf1b
Tnfrsf18
Pdcd1
Itgae
Itgb8
Entpd1
Nt5e
Lrrc32
Ikzf2
Ikzf4
Il10
Tgfb1
Ebi3
Pde3b
Cd40lg
Tcf7
Lef1
Ccr2
Ccr4
Ccr8
Il2
Il4
Ifng
2.0
1.0
0
-1.0
-2.0
z-score
Control
dbcAMP+Dex.
F
Down
Up
2e-10 Monocarboxylic acid metabolic process
1e-13 Small molecule metabolic process
2e-10 ATP generation from ADP
2e-10 Glycolytic process
1e-10 Nucleoside diphosphate phosphorylation
3e-12 Ribonucleoside diphosphate metabolic process
2e-12 Purine nucleoside diphosphate metabolic process
2e-11 ADP metabolic process
6e-11 Pyruvate metabolic process
1e-10 Cellular response to hypoxia
1e-11 Cellular response to decreased oxygen levels
1e-12 Cellular response to oxygen levels
1e-12 Response to hypoxia
4e-13 Response to decreased oxygen levels
2e-14 Response to oxygen levels
3e-04 Negative regulation of response to stimulus
2e-04 Cell surface receptor signaling pathway
3e-04 Regulation of multicellular organismal development
4e-06 Regulation of multicellular organismal process
3e-04 Localization of cell
3e-04 Cell motility
3e-04 Cell death
1e-04 Regulation of cell death
3e-05 Hematopoietic or lymphoid organ development
3e-05 Hemopoiesis
3e-05 Immune system development
1e-05 Leukocyte differentiation
3e-04 Mononuclear cell differentiation
4e-04 Immune response
4e-06 Immune system process
G
Slc2a1 TPM
Hk2 TPM
Gapdh TPM
Satb1 TPM
Pdcd1 TPM
Ctla4 TPM
Control
dbcAMP+Dex.
H
Cont
Vit.C
I
% FoxP3+ cells
CD25 MFI
GITR MFI
CTLA4 MFI
Bcl2 MFI
Cont
Vit.C

**Fig. S3. Effects of cAMP analog, dexamethasone and ascorbate on S/F-iTreg function.**

(**A**) Effects of dibutyryl-cAMP (dbcAMP) and dexamethasone on C-iTregs and S/F-iTregs. Percentages of Foxp3$^{+}$. IL-10$^{+}$ cells and Ki-67$^{+}$ cells assessed by flow cytometry (n=3). (**B** and **C**) MFI of CTLA-4$^{+}$ cells (B) and percentages of cytokine producing cells (C) in control or dbcAMP/dexamethasone-treated S/F-iTregs assessed by flow cytometry (n=3). (**D**) Suppressive activity of Treg populations. Proliferation of Treg-cocultured CFSE-labelled responder CD4$^{+}$ T cells (Tresp) in the presence of anti-CD3 and APCs was assessed and expressed as percentages compared with proliferation of Tresp cells alone as 100% (n=3). (**E-G**) RNA-sequencing analysis of dbcAMP/dexamethasone-treated S/F-iTregs. Gene expression pattern of Treg-related molecules (D), enrichment analysis of differential expressed genes (DEGs) (E) and gene expression levels (TPM) of typical DEGs (F) (n=3). (**H**) DNA hypomethylation of *Foxp3* CNS2 region in ascorbate-treated or non-treated S/F-iTregs assessed by bisulfite sequencing. Filled circle; methylated CpG, unfilled circle; demethylated CpG. Representative of 3 independent experiments. (**I**) Percentages and MFIs of Foxp3$^{+}$, CD25$^{+}$, GITR$^{+}$, CTLA-4$^{+}$ and Bcl2$^{+}$ cells assessed by flow cytometry (n=3). Vertical bars denote mean$\pm$SD in A-C, G, and I. Paired t-test for statistical analysis in B, C, G and I (* $p<0.05$, ** $p<0.01$). ANOVA followed by Dunnett's test for statistical analysis in A (* $p<0.05$ vs. control).

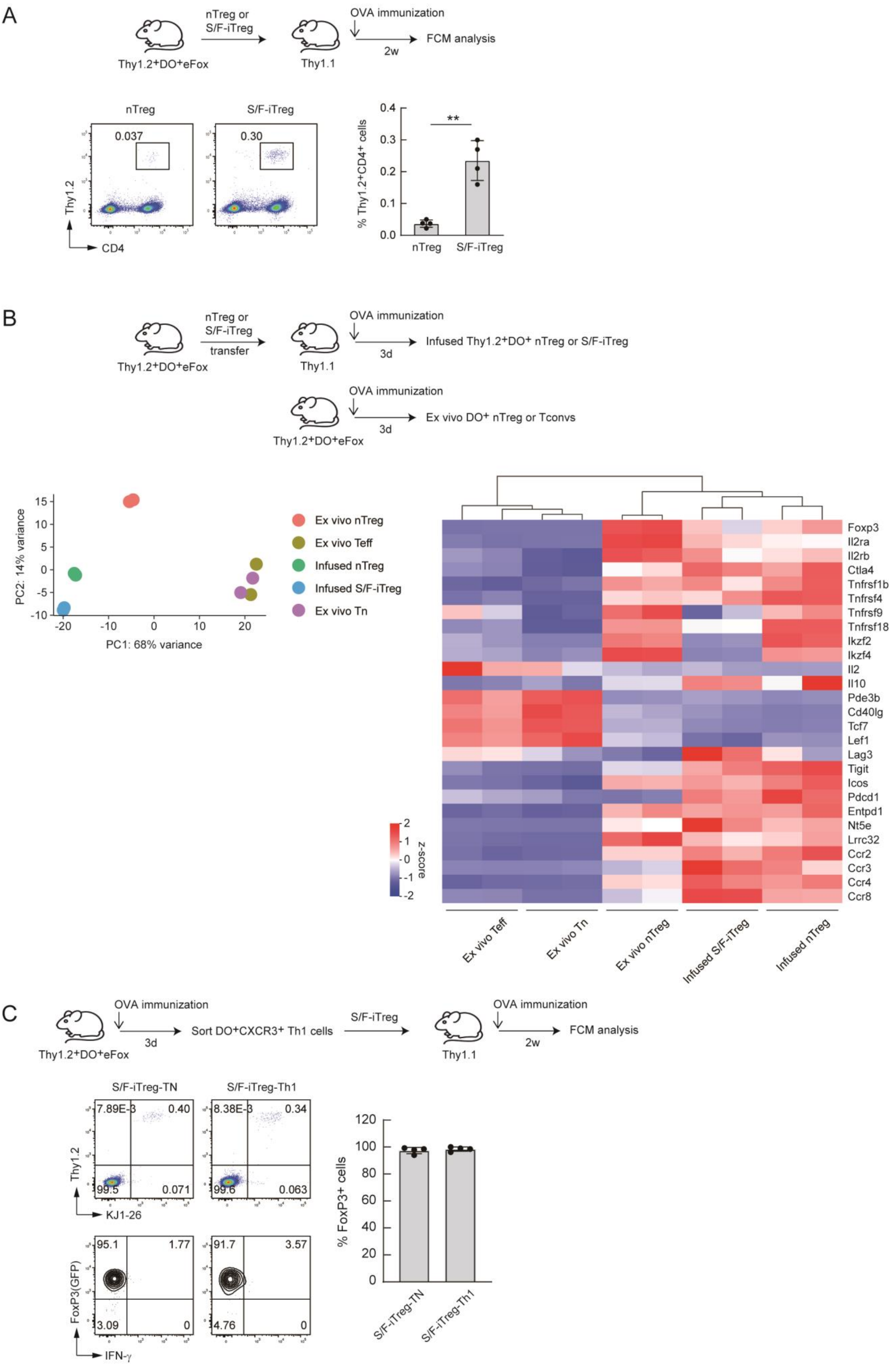
A
nTreg or
S/F-iTreg
OVA immunization
FCM analysis
2w
Thy1.2+DO+eFox
Thy1.1
nTreg
S/F-iTreg
0.037
0.30
Thy1.2
CD4
% Thy1.2+CD4+ cells
**
0.4
0.3
0.2
0.1
0.0
nTreg
S/F-iTreg
B
nTreg or
S/F-iTreg
transfer
OVA immunization
Infused Thy1.2+DO+ nTreg or S/F-iTreg
3d
Thy1.2+DO+eFox
Thy1.1
OVA immunization
Ex vivo DO+ nTreg or Tconvs
3d
Thy1.2+DO+eFox
PC2: 14% variance
PC1: 68% variance
15
10
5
0
-5
-10
-20
-10
0
10
20
Ex vivo nTreg
Ex vivo Teff
Infused nTreg
Infused S/F-iTreg
Ex vivo Tn
z-score
2
1
0
-1
-2
Foxp3
Il2ra
Il2rb
Ctla4
Tnfrsf1b
Tnfrsf4
Tnfrsf9
Tnfrsf18
Ikzf2
Ikzf4
Il2
Il10
Pde3b
Cd40lg
Tcf7
Lef1
Lag3
Tigit
Icos
Pdcd1
Entpd1
Nt5e
Lrrc32
Ccr2
Ccr3
Ccr4
Ccr8
Ex vivo Teff
Ex vivo Tn
Ex vivo nTreg
Infused S/F-iTreg
Infused nTreg
C
OVA immunization
3d
Sort DO+CXCR3+ Th1 cells
S/F-iTreg
OVA immunization
2w
FCM analysis
Thy1.2+DO+eFox
Thy1.1
S/F-iTreg-TN
S/F-iTreg-Th1
7.89E-3
0.40
99.5
0.071
8.38E-3
0.34
99.6
0.063
Thy1.2
KJ1-26
95.1
1.77
3.09
0
91.7
3.57
4.76
0
FoxP3(GFP)
IFN-γ
% FoxP3+ cells
120
100
80
60
40
20
0
S/F-iTreg-TN
S/F-iTreg-Th1

**Fig. S4.** ***In vivo*** **expansion/survival of OVA-specific nTregs and S/F-iTregs upon OVA stimulation.**

(**A**) *In vivo* persistency of DO11.10 nTregs and S/F-iTregs. Thy1.2 Tregs were transferred into Thy1.1 WT mice and immunized with OVA. Percentages of $Foxp3^+$ cells and $Thy1.2^+$ cells in dLN assessed by flow cytometry 2 weeks later. Vertical bars denote mean$\pm$SD (n=4). Unpaired t-test for statistical analysis (** p<0.01). (**B**) RNA-sequencing analysis of *in vivo* transferred S/F-iTregs. PCA and the analysis of the gene expression patterns of Treg-related molecules on $DO11.10^+$ S/F-iTregs and nTregs after transfer into WT mice and OVA immunization (shown as infused S/F-iTreg or nTreg, respectively), and freshly isolated nTregs, naïve T convs and effector T convs after immunization of DO11.10 mice (shown as ex vivo nTreg, Tn, and Teff, respectively) (n=2). (**C**) *In vivo* stability of Th1-derived S/F-iTregs. DO11.10 TCR transgenic mice were immunized with OVA; and $CXCR3^+$ or $CXCR3^-$ cells collected from draining LNs were subjected to S/F-iTreg conversion as shown in Fig. 3. S/F-iTreg-Th1 and S/F-iTreg-TN thus prepared were transferred into congenic mice and OVA immunized, and examined 2 weeks later for the expression of FoxP3 and IFN-γ by FCM (n=4).

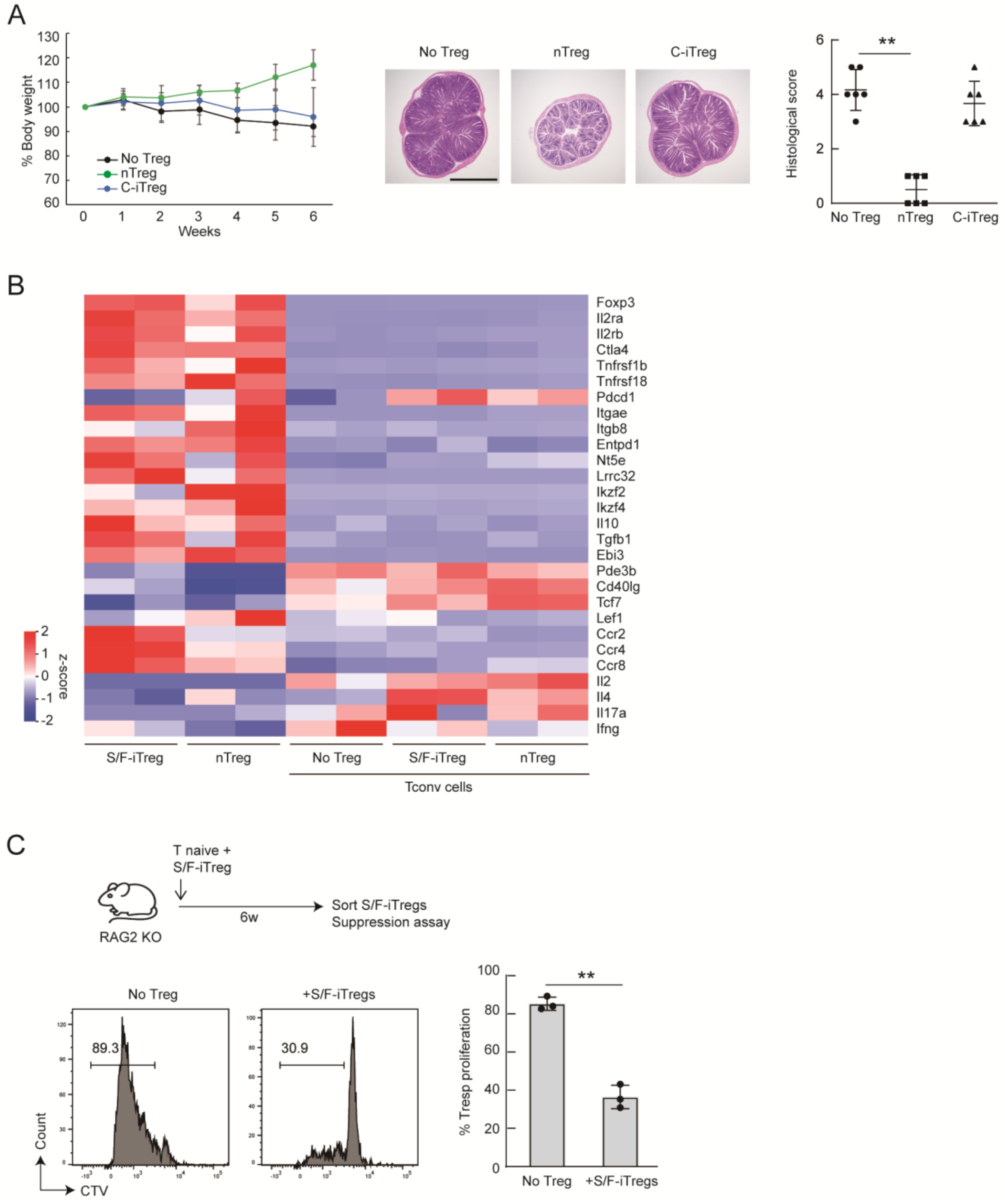
A
% Body weight
Weeks
No Treg
nTreg
C-iTreg
Histological score
**
B
Foxp3
Il2ra
Il2rb
Ctla4
Tnfrsf1b
Tnfrsf18
Pdcd1
Itgae
Itgb8
Entpd1
Nt5e
Lrrc32
Ikzf2
Ikzf4
Il10
Tgfb1
Ebi3
Pde3b
Cd40lg
Tcf7
Lef1
Ccr2
Ccr4
Ccr8
Il2
Il4
Il17a
Ifng
z-score
S/F-iTreg
nTreg
No Treg
S/F-iTreg
nTreg
Tconv cells
C
T naive +
S/F-iTreg
RAG2 KO
6w
Sort S/F-iTregs
Suppression assay
No Treg
+S/F-iTregs
89.3
30.9
Count
CTV
% Tresp proliferation
No Treg
+S/F-iTregs

**Fig. S5. *In vivo* suppressive activity of nTregs and C-iTregs on colitis.**

(**A**) Colitis development in CD45.1/RAG2 KO mice by transfer of Thy1.1$^+$CD4$^+$CD45RB$^{hi}$ naïve T cells with or without Thy1.2$^+$ nTregs or C-iTregs. Relative body weight expressed as percentages of the weight before cell transfer. Histological analysis of large intestine at the end point. Bar = 1 mm. Vertical bars denote mean$\pm$SD (n=6). (**B**) RNA-sequencing analysis of *in vivo* transferred S/F-iTregs, nTregs, and Tconvs that were transferred with S/F-iTregs, nTregs, or alone (no Tregs). The gene expression patterns of Treg-related molecules 6 weeks after cell transfer (n=2). (**C**) *In vitro* suppressive activity of infused S/F-iTregs. Thy-1.1$^+$ S/F-iTregs were transferred with Thy-1.2 naïve CD4$^+$ T cells to RAG2KO mice, which were analyzed 6 weeks later for suppressive activity. Thy-1.1$^+$ CD4$^+$ T cells were assessed for suppressing CTV-labelled CD4$^+$ T cells from BALB/c mice in the presence of BALB/c APCs (n=3). ANOVA followed by Turkey's post hoc test in A and paired t-test in C for statistical analysis (** $p<0.01$). Bar = 1 mm.

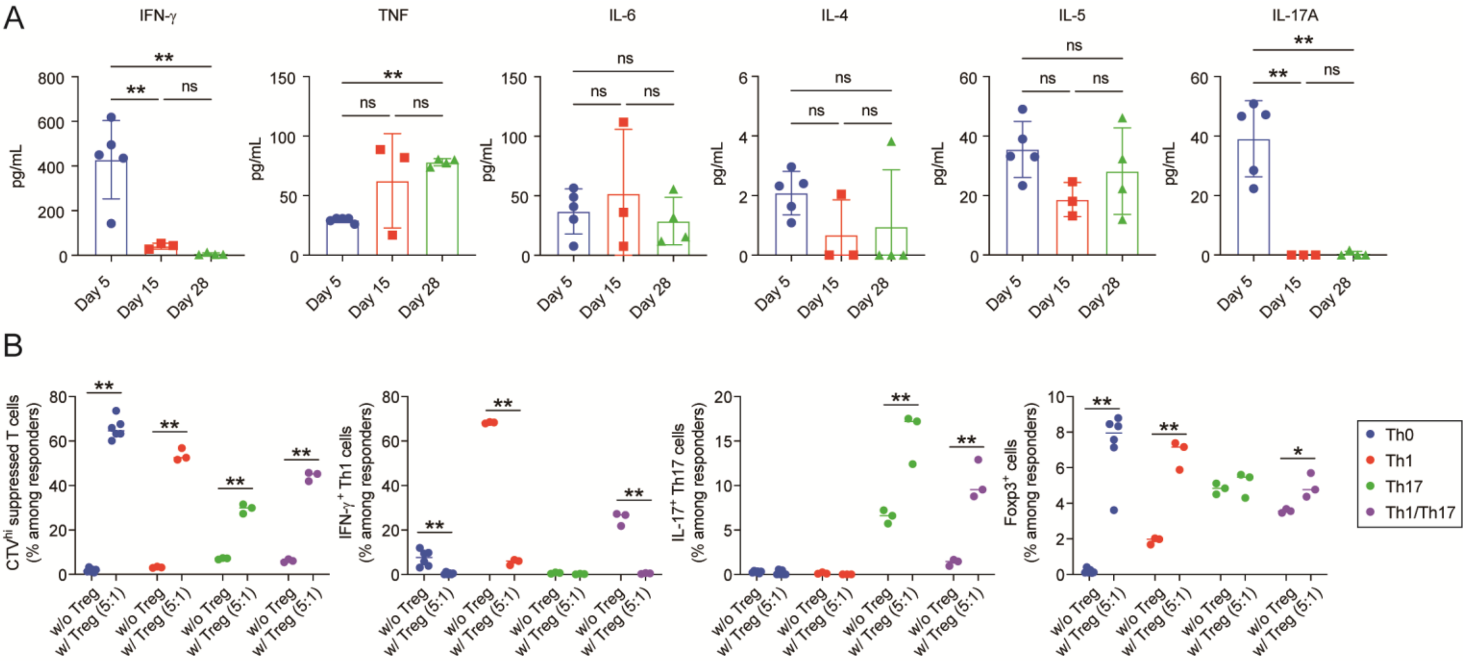

A
IFN-γ
TNF
IL-6
IL-4
IL-5
IL-17A
pg/mL
Day 5
Day 15
Day 28
ns
**
B
CTV^hi suppressed T cells (% among responders)
IFN-γ+ Th1 cells (% among responders)
IL-17+ Th17 cells (% among responders)
Foxp3+ cells (% among responders)
w/o Treg
w/ Treg (5:1)
Th0
Th1
Th17
Th1/Th17

**Fig. S6. Suppressive function of S/F-iTregs to control inflammatory cytokine production.**

(**A**) Kinetics of serum cytokine level in 5, 15, and 28 days after cell transfer with allogenic BM, splenocytes, and S/F-iTregs, in murine allogenic GVHD model shown in Figure 6A. Serum samples collected from inferior vena cava were measured with Cytometry Beads Array (CBA) (n=3-5). Data at day 5 are adopted from Figure 6F. (**B**) Suppressive function of S/F-iTregs in the control of Th differentiation. Responder naive $CD4^+$ T cells were labeled with CTV followed by co-culture with irradiated T cell-depleted splenocytes with or without S/F-iTregs (5 to 1 ratio to responder) for 4 days in the indicated Th conditions. Th0; anti-CD3 Ab alone, Th1; anti-CD3 Ab and IL-12, Th17; anti-CD3 Ab, IL-6 and TGF-β, Th1/Th17; anti-CD3 Ab, IL-12, IL-6 and TGF-β. The ratio of $CTV^{hi}$ non-proliferated cells, $Foxp3^-IFN\text{-}\gamma^+$ Th1 cells, $Foxp3^-IL\text{-}17^+$ Th17 cells, $Foxp3^+$ cells among responder T cells were analyzed with flowcytometry. Data show results from 3-6 replicates. ANOVA followed by Turkey's post hoc test in A and paired t-test in B for statistical analysis (* $p<0.05$, ** $p<0.01$).

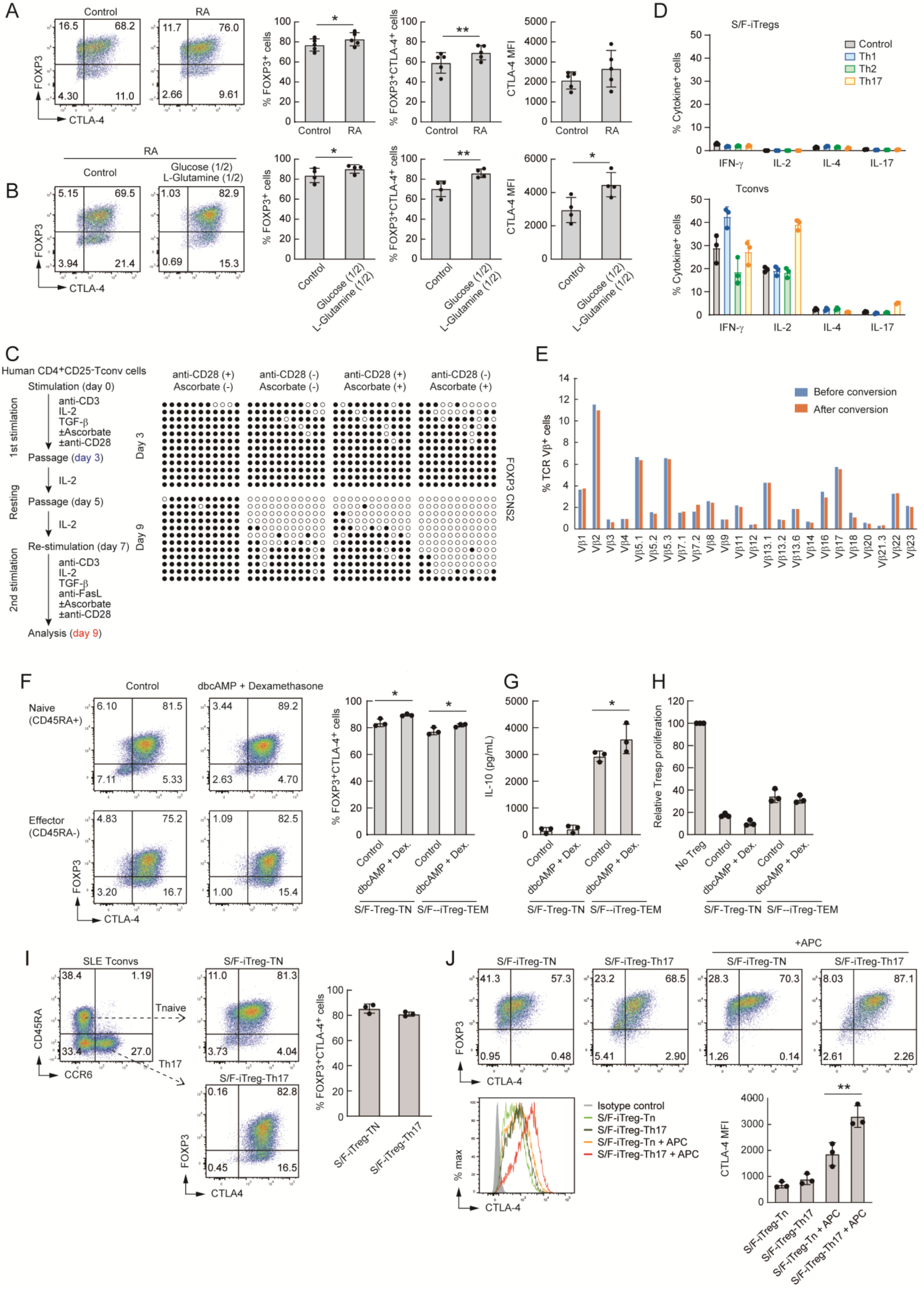
A
Control
RA
16.5
68.2
4.30
11.0
11.7
76.0
2.66
9.61
FOXP3
CTLA-4
% FOXP3+ cells
% FOXP3+CTLA-4+ cells
CTLA-4 MFI
Control
RA
*
**
B
RA
Control
Glucose (1/2)
L-Glutamine (1/2)
5.15
69.5
3.94
21.4
1.03
82.9
0.69
15.3
C
Human CD4+CD25-Tconv cells
Stimulation (day 0)
1st stimlation
anti-CD3
IL-2
TGF-β
±Ascorbate
±anti-CD28
Passage (day 3)
Resting
IL-2
Passage (day 5)
Re-stimulation (day 7)
2nd stimlation
anti-FasL
Analysis (day 9)
anti-CD28 (+) Ascorbate (-)
anti-CD28 (-) Ascorbate (-)
anti-CD28 (+) Ascorbate (+)
anti-CD28 (-) Ascorbate (+)
Day 3
Day 9
FOXP3 CNS2
D
S/F-iTregs
Tconvs
% Cytokine+ cells
Control
Th1
Th2
Th17
IFN-γ
IL-2
IL-4
IL-17
E
% TCR Vβ+ cells
Before conversion
After conversion
Vβ1
Vβ2
Vβ3
Vβ4
Vβ5.1
Vβ5.2
Vβ5.3
Vβ7.1
Vβ7.2
Vβ8
Vβ9
Vβ11
Vβ12
Vβ13.1
Vβ13.2
Vβ13.6
Vβ14
Vβ16
Vβ17
Vβ18
Vβ20
Vβ21.3
Vβ22
Vβ23
F
Control
dbcAMP + Dexamethasone
Naive (CD45RA+)
Effector (CD45RA-)
6.10
81.5
7.11
5.33
3.44
89.2
2.63
4.70
4.83
75.2
3.20
16.7
1.09
82.5
1.00
15.4
% FOXP3+CTLA-4+ cells
dbcAMP + Dex.
S/F-Treg-TN
S/F--iTreg-TEM
G
IL-10 (pg/mL)
H
Relative Tresp proliferation
No Treg
I
SLE Tconvs
38.4
1.19
33.4
27.0
CD45RA
CCR6
Tnaive
Th17
S/F-iTreg-TN
11.0
81.3
3.73
4.04
S/F-iTreg-Th17
0.16
82.8
0.45
16.5
CTLA4
J
+APC
S/F-iTreg-TN
S/F-iTreg-Th17
41.3
57.3
0.95
0.48
23.2
68.5
5.41
2.90
28.3
70.3
1.26
0.14
8.03
87.1
2.61
2.26
% max
Isotype control
S/F-iTreg-Tn
S/F-iTreg-Th17
S/F-iTreg-Tn + APC
S/F-iTreg-Th17 + APC

**Fig. S7. Effects of retinoic acid, limitation of glucose and L-glutamine, cAMP analog and dexamethasone on human S/F-iTreg induction.**

(**A**) Effects of retinoic acid (RA) on human S/F-iTregs. Percentages and MFI of FoxP3$^{+}$ cells and CTLA-4$^{+}$ cells assessed by flow cytometry (n=5). (**B**) Effects of reduced glucose and L-glutamine concentrations in the culture media. Percentages and MFI of FoxP3$^{+}$ cells and CTLA-4$^{+}$ cells assessed by flow cytometry (n=4). (**C**) DNA hypomethylation of the FOXP3 CNS2 region assessed by bisulfite sequencing. Filled circle; methylated CpG, unfilled circle; demethylated CpG. Representative of at least 2 independent experiments. (**D**) Cytokine production in re-stimulated S/F-iTregs and activated Tconvs assessed by flow cytometry (n=3). (**E**) TCR repertoire before and after S/F-iTreg conversion. The percentages of T cells expressing particular TCR Vβ subfamilies were analyzed by flow cytometry before and after S/F-iTreg conversion. Data is representative of three independent donors. (**F-H**) Effects of dbcAMP (20 μM) and Dexamethasone (10 nM) on human naïve- or effector/memory-derived S/F-iTregs. Percentages of FoxP3$^{+}$ cells and CTLA-4$^{+}$ cells assessed by flow cytometry (F), IL-10 production assessed by ELISA (G), and suppressive activity (H) (n=3). (**I**) S/F-iTreg induction from CCR6$^{+}$ CD45RA$^{-}$ effector/memory CD4$^{+}$ T cells of SLE patients. Percentages of FoxP3$^{+}$ and CTLA-4$^{+}$ cells assessed by flow cytometry (n=3). (**J**) Re-stimulation assay of S/F-iTregs with autologous monocyte DCs. MFI of CTLA-4 assessed by flow cytometry (n=3). Vertical bars denote mean$\pm$SD in A, B, and D, F-H. Paired t-test for statistical analysis in A and B (* $p<0.05$, ** $p<0.01$). ANOVA followed by SNK test for statistical significance for F, G and J (* $p<0.05$, ** $p<0.01$).

**Table S1. List of antibodies.**

| Antibody | Clone | Application | Vendor | Catalog# |
|---|---|---|---|---|
| anti-human CD152 (CTLA-4) | BNI3 | FCM | BD | BV421(562743) |
| anti-human CD178 (FasL) | NOK-1 | Cell culture | BioLegend | 306408 |
| anti-human CD196 (CCR6) | G034E3 | FCM | BioLegend | PerCP-Cy5.5(353405) |
| anti-human CD2 | RPA-2.10 | FCM | BioLegend | PE-Cy7(300222) |
| anti-human CD25 | BC96 | FCM | BioLegend | FITC(302604)<br>BV421(302630) |
| anti-human CD28 | 15E8 | Cell culture | Milteny | 170-076-117 |
| anti-human CD3 | OKT3 | Cell culture | Milteny | 170-076-116 |
| anti-human CD4 | RPA-T4 | FCM | Invitrogen | APC(17-0049-42) |
| anti-human CD45RA | HI100 | FCM | BD | FITC(555488) |
| anti-human FOXP3 | 236A/E7 | FCM | Invitrogen | PE(12-4777-42) |
| anti-human IFNγ | 4S.B3 | FCM | BioLegend | BV421(502532) |
| anti-human IL-2 | MQ1-17H12 | FCM | BD | FITC(554565) |
| anti-human IL-4 | 8D4-8 | FCM | BD | APC(560671) |
| anti-human IL-17A | eBio64DEC17 | FCM | Invitrogen | PE(12-7179-42) |
| anti-human/mouse CD44 | IM7 | FCM | Invitrogen | APC(17-0441-83) |
| anti-human/mouse GATA3 | TWAJ | FCM | Invitrogen | eFluor660(50-9966-42) |
| anti-human/mouse Helios | 22F6 | FCM | BioLegend | APC(137218) |
| anti-human/mouse Histone H3K27ac | Polyclonal | ChIP | GeneTex | Unconjugated(GTX60815) |
| anti-mouse Bcl2 | BCL/10C4 | FCM | BioLegend | PE(633508) |
| anti-mouse CD120b (TNFR2) | TR75-89 | FCM | BioLegend | PE(113405) |
| anti-mouse CD134 (OX40) | OX-86 | FCM | BioLegend | BV421(119411) |

| | | | | |
|---|---|---|---|---|
| anti-mouse CD152 (CTLA-4) | UC10-4B9 | FCM | BioLegend | BV421(106312) |
| anti-mouse CD16/32 | 93 | FCM | BioLegend | Unconjugated(101302) |
| anti-mouse CD178 (FasL) | MFL3 | Cell culture | BioLegend | 106612 |
| anti-mouse CD183 (CXCR3) | CXCR3-173 | FCM | BioLegend | PE(126506) |
| anti-mouse CD193(CCR3) | 83103 | FCM | BD | APC(557974) |
| anti-mouse CD194(CCR4) | 2G12 | FCM | BioLegend | BV421(131217) |
| anti-mouse CD196(CCR6) | 140706 | FCM | BD | Alexa Fluor647(557976) |
| anti-mouse CD223 (Lag3) | C9B7W | FCM | BD | PE(552380) |
| anti-mouse CD25 | PC61<br>PC61.5 | FCM | BioLegend<br>Invitrogen | BV421(102034)<br>APC(17-0251-82) |
| anti-mouse CD278 (ICOS) | C398.4A | FCM | BioLegend | PE-Cy7(313520) |
| anti-mouse CD28 | 37.51 | Stimulation | BD | 553294 |
| anti-mouse CD357 (GITR) | DTA-1 | FCM | BD | PE-Cy7(558140) |
| anti-mouse CD366 (Tim3) | RMT3-23 | FCM | Invitrogen | PE(12-5870-81) |
| anti-mouse CD38 | 90 | FCM | Invitrogen | APC(17-0381-82) |
| anti-mouse CD3ε | 145-2C11 | Stimulation | BD | 553057 |
| anti-mouse CD4 | RM4-5 | FCM | BioLegend | BV605(100548)<br>BV711(100550) |
| anti-mouse CD45.1 | A20 | FCM | BioLegend | PE-Cy7(110780) |
| anti-mouse CD45.2 | 104 | FCM | BioLegend<br>BD | BV421(109832)<br>PerCP-Cy5.5(552950) |
| anti-mouse CD62L | MEL-14 | FCM | BD | PerCP-Cy5.5(560513) |
| anti-mouse CD8a | 53-6.7 | FCM | BioLegend | PerCP-Cy5.5(100734) |

| | | | | |
|---|---|---|---|---|
| anti-mouse CD90.1 (Thy1.1) | OX-7 | FCM | BioLegend | BV421(202529) |
| anti-mouse CD90.2 (Thy1.2) | 53-2.1 | FCM | BD | APC(553007),<br>PE(553006) |
| anti-mouse DO11.10 TCR | KJ1-26 | FCM | BD<br>Invitrogen | PE(551772)<br>APC(17-5808-80) |
| anti-mouse GARP | YGIC86 | FCM | Invitrogen | PE(12-9891-82) |
| anti-mouse GM-CSF | MP1-22E9 | FCM | BD | PE(554406) |
| anti-mouse H-2Kb | AP6-88.5 | FCM | BioLegend | Biotin(116504)<br>Alexa Fluor488(116510)<br>PE(116508) |
| anti-mouse H-2Kd | SF1-1.1 | FCM | BioLegend | APC-cy7(116630) |
| anti-mouse IFNγ | XMG1.2 | FCM | BioLegend<br>BD<br>Invitrogen | BV421(505830)<br>Alexa Fluor647(557735)<br>PE(12-7311-82) |
| anti-mouse IL-10 | JES5-16E3 | FCM | BioLegend | PE-Cy7(505026)<br>Alexa Fluor647(505014) |
| anti-mouse IL-17A | TC11-18H10.1 | FCM | BD | PE(559502) |
| anti-mouse IL-2 | JES6-5H4 | FCM | BioLegend | BV421(503825) |
| anti-mouse IL-4 | 11B11 | FCM | BD | PE(554435) |
| anti-mouse Ki-67 | SolA15 | FCM | Invitrogen | PE-Cy7(25-5698-82) |
| anti-mouse LAP (TGF-β1) | TW7-16B4 | FCM | BioLegend | BV421(141407) |
| anti-mouse RORγt | Q31-378 | FCM | BD | PE(562607) |
| anti-mouse T-bet | 4B10 | FCM | BioLegend | BV421(644816) |
| anti-mouse TCR-beta | H57-591 | FCM | BioLegend | BV785(109249) |
| anti-mouse TIGIT | 1G9 | FCM | BioLegend | APC(142105) |
| anti-mouse TNF | MP6-XT22 | FCM | BD | PE(554419) |
| anti-mouse/rat FoxP3 | FJK-16s | FCM | Invitrogen | |
| Streptavidin | | FCM | BD | PE(554061) |
